# Metabolic impact of trans 10, cis 12-conjugated linoleic acid on pai transgenic mice

**DOI:** 10.1101/2023.01.31.526123

**Authors:** Yu Rao, Shi-Li Li, Mei-Juan Li, Bao-Zhu Wang, Yang-Yang Wang, Lu-Wen Liang, Shuai Yu, Zong-Ping Liu, Sheng Cui, Ke-Mian Gou

## Abstract

*Trans* 10, *cis* 12-conjugated linoleic acid (t10c12-CLA) from ruminant-derived foodstuffs can induce body fat loss in mammals after oral administration, while its mechanism on fat reduction has yet to be clarified fully until now. In the current study, a transgenic mouse that produced t10c12-CLA had been generated by inserting the Propionibacterium acnes isomerase (Pai) expression cassette into the Rosa26 locus, and its male offspring were used to decipher an irreversible long-term impact of t10c12-CLA on health and its mechanism of action. Compared to their wild-type C57BL/6J littermates, comprehensive phenotype profiling of biallelic pai/pai mice indicated that white fat was decreased while brown fat was increased reversely; meanwhile, more heat was released and the central activities were reduced. Besides decreased plasma triglycerides in both pai genotypes and increased serum FGF21 in pai/wt mice, RNA and protein analysis revealed that the fatty acid oxidation and thermogenesis capacity of brown adipose tissues were elevated via CPT1B and UCP1/2 over-expression. The results indicate that the t10c12-CLA-induced fat loss might be caused by the excess FGF21 and the increased mass and extra thermogenesis of brown adipose tissue in transgenic mice.

## 1. Introduction

*Trans* 10, *cis* 12-conjugated linoleic acid (t10c12-CLA), one of the isomers of linoleic acid (18:2n-6, LA), is naturally present in ruminant-derived foodstuffs. A person is estimated to take about 9∼20 mg of t10c12-CLA from foodstuffs daily [1]. Since the effects of t10c12-CLA on body fat loss in mice [2] and milk fat depression in cows [3] were reported, numerous studies tried to explore its mechanism on lipid metabolism. Some studies revealed that t10c12-CLA induces fat reduction by reducing food intake, modulating lipid metabolism, increasing energy expenditure, and/or browning white adipose tissue (WAT) [4-6]. At times, some of these reports showed contradictory results. Thus, the mechanism of action of t10c12-CLA on body fat reduction has yet to be clarified fully until now.

Meanwhile, other studies also indicated that t10c12-CLA influenced the metabolism or physiological roles of some critical fatty acids (FAs), such as palmitic, stearic, oleic, linoleic, arachidonic, docosahexaenoic acids and/or their derived leukotrienes signals/hormonal molecules via affecting their contents in various tissues [7-10] depending on the level of dietary fat and the degree of host obesity [9,11]. Furthermore, these studies in mice fed with a t10c12-CLA diet revealed approximately 0.1-1.5% of t10c12-CLA in the brain and eye [12], milk [8], adipose tissues [7], liver [7,9], heart, spleen, kidney, muscle, and serum [9], suggesting that t10c12-CLA might have a systemic impact on the body. In fact, besides anti-obesity, a few studies also showed the connections between t10c12-CLA and hepatic steatosis [5,13], pro-inflammatory effects [13], and even intestinal microbiota in mice [14], suggesting the potential risk of using t10c12-CLA.

T10c12-CLA is present at very low concentrations in nature, while most t10c12-CLA studies or applications used a chemically CLA mixture prepared by industrial isomerisation. CLA mixture usually contains approximately equal amounts of t10c12-CLA and *cis* 9, *trans* 11-CLA (c9t11-CLA), typically 80–95% of total CLA, and other minor isomers, such as *trans*, *trans* isomers [6]. To reveal the entire and exact t10c12-CLA’s impact that might be offset or masked by c9t11-CLA [6] or trans fatty acids [15] in the CLA mixtures, we seek the possibility of producing t10c12-CLA by an animal itself so we might investigate its long-term impact and mechanism of action in a novel way.

Based on the reports that *Propionibacterium acnes* isomerase (PAI) can successfully convert LA into t10c12-CLA in recombinant lactic acid bacteria [7,9] and transgenic plants [16], We produced t10c12-CLA efficiently in murine 3T3 cells [17]. Then we established a pai-transgenic mouse previously [18]; unfortunately, the pai-expressing cassette was randomly integrated into the Myh11 gene in this mouse, resulting in the death of homozygous mice after birth due to gastric inflation and bladder without urinating ability [19].

In the present study, we produced a novel transgenic founder in which the pai-expressing cassette was knocked into the Rosa26 locus by the CRISPR-Cas9 technique. The results indicated that the pai/pai mice exhibited reduced white fat and lowered triglycerides (TGs) in the blood but had no changes in hormones involved in energy intake, stress response, inflammation, and fatty liver, suggesting long-term exposure to t10c12-CLA is safe, at least in male mammals. Furthermore, we first revealed that increased FGF21 in the blood and more heat released by brown fat resulted in white fat reduction in pai mice. The results shed light on the mechanism of action of t10c12-CLA. Moreover, aberrant expression of the hypothalamic genes suggests that t10c12-CLA could have other potential impacts on health. Thus, the use of t10c12-CLA in anti-obesity practice or clinic trials should be closely monitored to ensure proper and safe application.

## 2. Materials and Methods

### 2.1. Chemicals

Unless otherwise stated, all chemicals were purchased from Sigma (USA) or Sinopharm Chemical Reagent Co., Ltd. (Shanghai, China). Restriction endonucleases were purchased from New England Biolabs Inc (USA).

### 2.2. Vector construction

The pRosa-pai vector was constructed as follows. In brief, based on the nucleotide sequence of the Pai gene of *Propionibacterium acnes* (GenBank accession number AX062088), the codon-optimized Pai gene was synthesised, which added a 6 x His affinity tag sequence immediately upstream of the terminal codon. It has been confirmed that gene transfection with this vector in murine 3T3 cells led to t10c12-CLA production [17]. On this basis, the Pai expression vector was firstly generated after insertion into the *Eco*R I site of the pCAGGS vector (Gift from Dr Timothy J. Ley, The Washington University, St. Louis), in which the Pai gene was under the control of the cytomegalovirus enhancer and the chicken beta-actin promoter (CAG promoter). After two molecular cloning steps of enzymic modification, the CAG-pai expression cassette was inserted into the Pac I and Asc I sites of the pROSA26-PA backbone vector (Gift from Dr Frank Costantini, Addgene plasmid#21271; http://n2t.net/addgene:21271) to generate the pRosa-pai vector.

The pRosa-Cas9 vector was constructed as follows. Briefly, the CAG promoter sequence of Cas9 in pDG330 plasmid (Gift from Dr Paul Thomas; Addgene plasmid # 100898; http://n2t.net/addgene:100898) was replaced by the EF-1α promoter from the plasmid pEF.myc.ER-E2-Crimson (Gift from Dr Benjamin Glick; Addgene plasmid # 38770; http://n2t.net/addgene:38770) followed by insertion of the sgRNAs sequences (5-actccagtctttctagaaga-3, from reference [20]) into Bbs I sites to generate the pRosa-Cas9 vector which included the Cas9 gene driven by EF-1α promoter and dual gRNA expression cassettes.

### 2.3. Pronuclear microinjection

Purified pRosa-Cas9 plasmid DNA at a final concentration of 10 ng/µl and the fragments of pRosa-pai plasmid after digestion by Pvu I and Sal I enzymes at a final concentration of 50∼100 ng/µl were mixed well and used for pronuclear co-microinjection immediately.

### 2.4. Mice

Mice were obtained from the Laboratory Animal Centre, Yangzhou University, China. The animal study protocol was approved by the Ethics Committee of Yangzhou University (protocol code NSFC2020-SYXY-20 and dated 25 March 2020). Mice were maintained in a light-controlled room (12L:12D, lights on at 0700 h) at a temperature of 22-23°C and were fed ad libitum with a standard diet containing 10% kcal% fat.

Embryos from superovulated D6B2F1 females mated with C57BL/6J male mice were used to produce the transgenic mice. The transgenic founders were backcrossed to C57BL/6J mice more than eight generations before analysis in the current study. The biallelic (pai/pai), monoallelic (pai/wt), and wild-type (wt) male offsprings from the pai/wt matings were used in this study.

### 2.5. Diets

New diets containing 10% kcal% fat were prepared per month under sterile conditions according to formula No. D12450H of OpenSource DIETS^TM^ (Research Diets Inc., NJ, USA). The formula D12450H and the sources of the food-grade ingredients for the diets are listed in Supplementary Table 1. Fatty acid compositions in diets were spot-checked by gas chromatography to examine the contents of linoleic acid and t10c12-CLA. One kilogram of diet contained approximately 20 g of LA, and no t10c12-CLA was detected in diet samples.

### 2.6. Transgene characterisation

Genomic DNA from the tail tissues was used for nucleic acid analysis. The presence of the pai transgene was assayed to amplify the 514-bp pai fragments (All primer sequences and annealing temperatures were listed in Supplementary Table 2). The same insertion site was assayed to amplify the 605-bp fragments spanned the insertional Rosa26 area. The PCR amplification was performed with 30 cycles at 94°C for 30 sec, 57°C for 30 sec, and 72°C for 40 sec.

Southern blot was performed according to the standard procedure. In brief, 10 µg of genomic DNA was first digested by the EcoR I enzyme for 12-16 h at 37°C and subjected to 1% agarose gel electrophoresis, then transferred onto the Hybond-N^+^ membrane (Amersham, UK). The membrane was subsequently hybridised with the DIG (Roche, Germany) labelled probes according to the manufacturer’s protocol, using alkaline phosphatase labelled anti-Dig antibody (Roche) and CDP-STAR solution to develop the photos.

### 2.7. Western blot

Western blot was performed according to the standard procedures. Briefly, total protein extracts from 20 mg tissues were quantified, and 20 µg protein extracts were separated with 10-15% SDS-PAGE gel and transferred to the PVDF membranes (Millipore ISEQ00010, Merck). Each membrane was blocked and slit into two parts. One was hybridised with the primary antibody to the protein of interest, and the other was hybridised with the antibody to beta-ACTIN (Proteintech Group, Inc, China, 20536-1-AP) or GAPDH (Abcam AB8245) followed by HPR-labelled goat antirabbit IgG (Santa Cruz Biotechnology, sc-2004). The chemiluminescent signal was developed using the SuperSignal™ West Femto substrate (ThermoFisher, USA). Blots were imaged for 5 s to 2 min and quantified using ImageJ software (NIH) and values were normalised to beta-ACTIN as a loading control. The primary antibodies included rabbit antibodies to His-tag (Proteintech Group, 10001-0-AP), CPT1B (Proteintech, 22170-1-AP), UCP2 (Proteintech, 11081-1-AP), AMPK ⍺1 (Abmart Shanghai Co., Ltd, China; T55326), pAMPKα (Thr172) (Abmart, TA3423), AMPK ⍺1 (phospho T183) + ⍺2 (phospho T172) (Abcam, AB133448), AMPK ⍺1 + ⍺2 (Abcam, ab207442), FASN (Abcam, AB128870), UCP1 (Abcam, ab234430), PPARγ (Wanleibio Ltd, China, WL01800), PGC1Α (Wanleibio, WL02123), or Phospho-HSL (Ser563, Cell Signaling Technology, #4139).

### 2.8. RNA analysis

Total RNA was extracted from tissues using the RNAiso Plus kit and treated with DNase I (TaKaRa, China). The purified RNA was used for first-strand cDNA synthesis, and reverse transcription was performed using an M-MLV reverse transcriptase with oligo-dT primers according to the manufacturer’s instructions (Promega, USA). For RT-PCR, the resulting cDNA was used for PCR amplification with the Pai-specific primers (Supplementary Table 2) that produced a 164-bp fragment. A 78-bp fragment of the Gapdh gene was amplified under the same conditions and used as an endogenous control. PCR amplification was performed as follows: 30 cycles of 94°C for 30 sec, 60°C for 30 sec, and 72°C for 30 sec.

All real-time PCRs were carried out in 96-well plates using ChamQ SYBR qPCR Master Mix kit in the ABI Prism 7500 Sequence Detection System (Applied Biosystems, USA) at 95°C, 2 min, one cycle; 95°C, 5 sec, 60°C, 32 sec, 40 cycles. Three replicates were included for each sample. The relative transcriptional level of a target gene was normalised to one of the endogenous gene expressions of Gapdh, ApoB, or 36B4 using the method of 2^-ΔΔCt^. All genes’ full names and these primer sequences are listed in Supplementary Table 3.

### 2.9. RNA sequencing

The RNA-seq was used to identify whole transcriptome differences in the brown adipose tissue (BAT) from wt (n=2) and pai/pai (n=3) mice at 12 weeks. Total RNAs were analysed by BGI Genomics company (China) using the DNBSEQ platform. Clean reads were aligned to *Mus musculus* genome GCF_000001635.26_GRCm38.p6, and the expression level was normalised as FPKM with gene annotation file. Differential expressional genes and functional enrichment for KEGG had plotted an online platform for data analysis and visualisation (http://www.bioinformatics.com.cn).

### 2.10. Gas chromatography

According to the modified method described by Jenkins [21], the procedure of two-step transesterification that utilised sodium methoxide followed by shorter methanolic HCl was used to methylate each organ/tissue homogenised by grinding in liquid nitrogen. Fatty acid methyl esters were separated on an HP-88 fused-silica capillary column (60 m X 0.25 mm i.d., 0.2-µm film thickness, J & W 112-88A7, Agilent Technologies, USA) and quantified using a fully automated 7890 Network GC System with a flame-ionization detector (Agilent). The program setting details followed Jenkins’ method [21]. C19:0 was used as the internal standard, and the peaks were identified by comparison with FA standards (Sigma, 47885U and O5632). The area percentage for all resolved peaks was analysed using GC ChemStation Software (Agilent).

### 2.11. Hepatic parameters measurements

Hepatic tissues (0.1 g) were homogenised and resuspended in PBS solution. The concentrations of total cholesterol (TC) and triglycerides (TGs) were respectively measured using the corresponding detecting kits (Meimian Industrial Co. Ltd., China).

### 2.12. Blood parameters measurements

Unless otherwise stated, all blood samples were collected from the 11∼15-week-old mice fed ad libitum. Fresh blood from the tail veins was used to measure the circulating glucose using a hand-held glucose monitor (Accu-Chek^®^ Performa Blood Glucose Meter, Roche). Heparin-treated blood from the tail vein was respectively used to measure the plasma TGs or high-density lipoprotein (HDL) using a hand-held cholesterol monitor (On-Call^®^ CCM-111 Blood Cholesterol System, Aikang Biotech Co. Ltd., China). Blood samples from the submandibular vein of conscious mice were collected at the first 4∼5 hrs of the light phase and used to measure the serum corticosterone using the Cort ELISA kit (Ruixin Biotech Co. Ltd., China). Serums from the orbital sinus vein of anaesthetic mice were used to measure the circulating TC, free fatty acids (FFA), prostaglandin E2 (PGE2) using the respective ELISA kits (Meimian), lactate dehydrogenase (Jiancheng Bioengineering Institute, Nanjing, China), as well as insulin, ghrelin, leptin, FGF21, interleukin-6 (IL-6), adrenaline glucagon tumour necrosis factor-alpha (TNFα), and C reactive protein (CRP) using the respective ELISA kits (Ruixin).

### 2.13. Intraperitoneally glucose and insulin tolerance tests

For the glucose tolerance test, each 11∼15-week-old mouse fasted overnight was intraperitoneally injected with D-glucose (2 g/kg body weight). For the insulin tolerance test, each mouse at 11-15 weeks fasted 4 h and was intraperitoneally injected with insulin (0.75 IU/kg body weight; Beyotime Biotech Inc, Shanghai, China). Blood glucose was measured in a drop collected from the tail vein prior to glucose or insulin injection and, 15, 30, 60, 90, and 120 min post glucose or insulin injection.

### 2.14. Histological analysis

Histological analysis was performed following the standard methods. Briefly, the tissues were fixed by immersion in 4% formaldehyde in PBS and then dehydrated before embedding. The 5-µm tissue sections (Leica CM1950) embedded with paraffin were dewaxed in xylene and rehydration. Histological details were examined using hematoxylin-eosin staining kits (Beijing Solarbio Science & Technology Co., Ltd., China). For red oil staining, 10-µm tissue slices embedded by Tissue-Tek^®^ O.C.T. Compound (Sakura, USA) were treated with a 60% isopropyl alcohol solution twice, and histological details were examined after staining with oil red O (Wuhan Servicebio Technology Co., Ltd., China). Stained slides were examined with an Olympus microscope equipped with a digital camera.

Each cellular cross-sectional area was estimated following the method described by Chen and Farese [22]. In detail, at least six slices from each mouse sample were randomly chosen for image analysis. Firstly, three photos of every slice were randomly taken from non-overlapping microscope fields under the 20 x objective. The cross-sectional area of each cell on an image was calculated one by one using ImageJ software, and the means of the top 100 cell areas per picture represented the value of a microscope sampling field. Finally, the means of at least 18 values per mouse sample (3 fields x 6 slices) were regard as this sample’s average cellular cross-sectional area for statistical analysis.

### 2.15. Metabolic cage measurements

Indirect calorimetry, heat production, and movement of 11-week-old mice were measured using the Automated Home Cage Phenotyping TSE PhenoMaster V4.5.3 system (TSE Systems Inc., Germany). The climate chamber was set to 22°C with a 12-h:12-h light-dark cycle (lights on at 0700 h). Mice were individually housed in plexiglass cages and fed ad libitum. VO_2_, VCO_2_, and food intake were measured every 39 min. After a 48 hrs acclimation period, the data were collected for the following 72 hrs, and the calculated lean mass was adjusted for all measurements.

### 2.16. Statistical analysis

All values are presented as mean ± SD. The statistical significance was assessed by unpaired student’s t-test or one-way ANOVA (Brown-Forsythe and Welch) among wt, pai/wt, and pai/pai mice. A value of p < 0.05 was considered statistically significant.

## 3. Results

### 3.1. Transgene analysis

The strategy to insert the pai-expressing cassette into the Rosa26 locus is shown in Figure 1A. PCR and Southern blot (Fig. 1 B-C) and DNA sequencing analysis demonstrated that the correct transgene was inserted into the Rosa26 locus in a founder mouse among 96 mice derived from injected embryos. RT-PCR and western blot analyses (Fig. 1 D-E) indicated that the pai gene was expressed in all tested tissues. Gas chromatography verified that the functional PAI enzyme successfully produced the t10c12-CLA (Fig. 1 F). Real-time PCR analysis (Fig. 1 G) revealed that the mRNA levels of Pai were quite different between biallelic pai/pai and monoallelic pai/wt mice; thus, two pai genotypes were analysed separately in the following tests.

**Figure 1.**
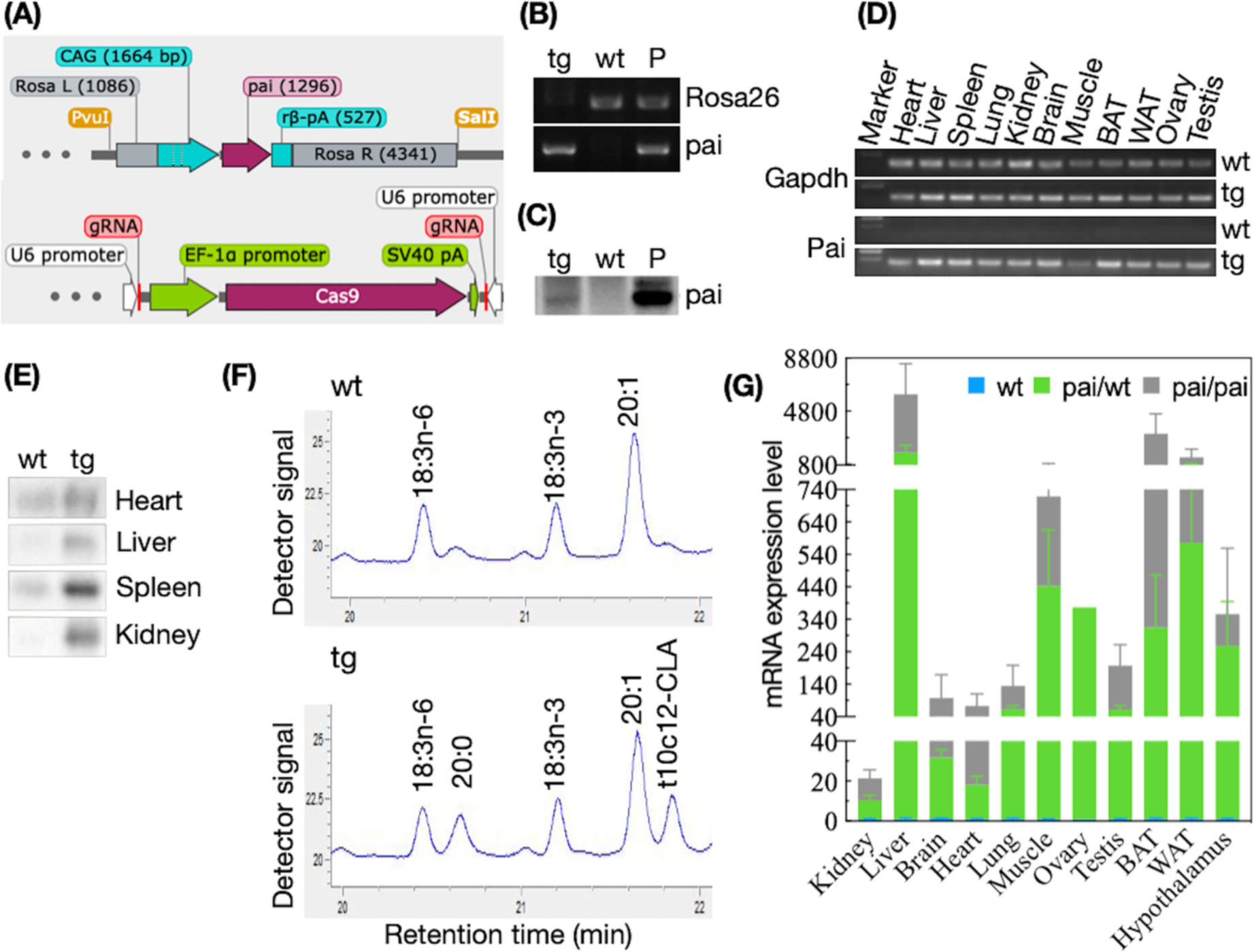
Transgene analysis. (A) Strategy for inserting the pai-expressing cassette into the Rosa26 locus by the CRISPR/Cas9 technique. The upper panel illustrates the recombined homological vector in which the codon-optimised Pai gene is driven by a CAG promoter and ended by a rabbit beta-globin polyA signal (rβ-pA). Rosa L and Rosa R represent the left and right sequences of the insertional site within the first intron of Rosa26, respectively. Linear DNA fragments digested with Pvu I and Sal I enzymes are used for comicroinjection. The down panel shows the CRISPR/Cas9 plasmid containing EF-1α-driven Cas9 and dual gRNA target sequences for co-microinjection. The numbers in brackets indicate the sequence length. (B) PCR and (C) Southern blot analysis of the genomic DNA samples from transgenic (tg), wild-type (wt) mice, and wt DNA mixed with a pRosa-Pai plasmid (P) shows that the Pai gene is inserted into the Rosa26 locus in the tg sample. PCR products are 514-bp fragments spanning the Pai gene and 605-bp fragments spanning the insertional site of the Rosa26 sequence. (D) RT-PCR analysis of RNAs shows that the Pai gene is expressed in all tg tissues, including brown (BAT) and white (WAT) adipose tissues, yet not in all wt samples. The amplified product of Pai is 164-bp fragments, and the Gapdh is 78-bp fragments as an internal control. (E) Western blot analysis reveals that PAI antigen existed in the tg heart, liver, spleen, and kidney, but not in wt samples. (F) The lipid profiles of partial gas chromatography traces show that the t10c12-CLA is produced in the kidney of the tg mouse compared to its wt littermate. (G) Real-time PCR analysis showed that the relative mRNA expression level of the Pai gene is quite different in each sample from biallelic pai/pai or monoallelic pai/wt mice (n=3), and no Pai mRNA is detected in samples of their wt littermates. Bars represent the mean ± SD after normalisation to the mRNA levels of Gapdh.

### 3.2. Fatty acids analysis

T10c12-CLA and FAs compositions in the hearts, livers, kidneys, tibialis anterior muscle, BAT, or WAT were analysed by gas chromatography. Compared to wt littermates, the contents (µg/g of tissue weight) of t10c12-CLA had increased in the kidneys (p = 0.004) and heart (p = 0.055) from pai/wt mice and WAT from pai/wt or pai/pai mice (p < 0.05; Table 1), not in the other examined tissues, suggesting that the t10c12-CLA is either produced in trace amounts or degraded quickly. Simultaneously, the quantities of LA substrates had decreased in the livers from both pai genotypes and increased in the pai/wt kidneys (p < 0.05; Table 1).

**Table 1.**
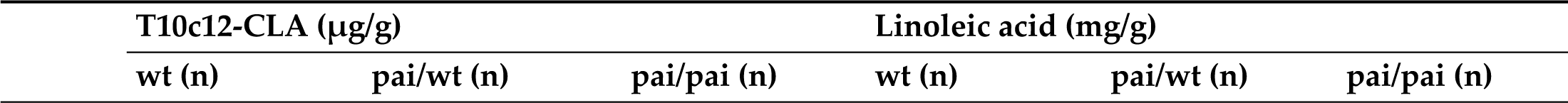
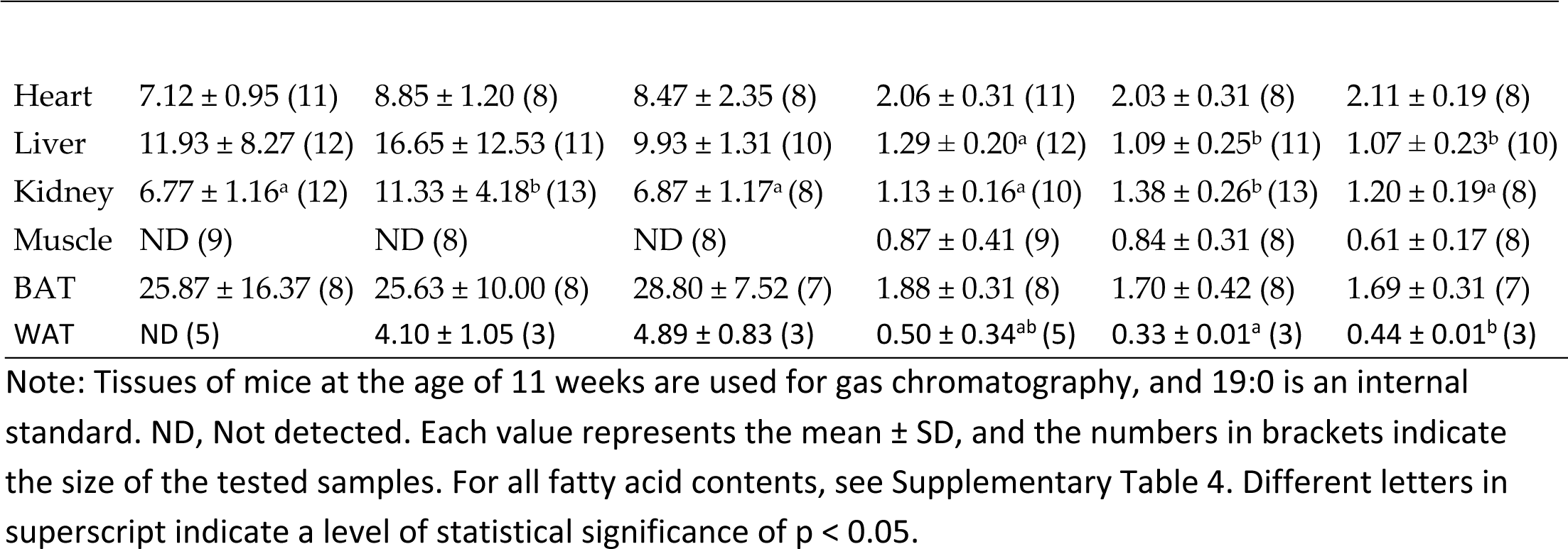
Contents of t10c12-CLA and linoleic acid in wt and pai mice.

| T10c12-CLA (μg/g) |  |  | Linoleic acid (mg/g) |  |  |
| --- | --- | --- | --- | --- | --- |
| wt (n) | pai/wt (n) | pai/pai (n) | wt (n) | pai/wt (n) | pai/pai (n) |

|  |  |  |  |  |  |  |
| --- | --- | --- | --- | --- | --- | --- |
| Heart | 7.12 ± 0.95 (11) | 8.85 ± 1.20 (8) | 8.47 ± 2.35 (8) | 2.06 ± 0.31 (11) | 2.03 ± 0.31 (8) | 2.11 ± 0.19 (8) |
| Liver | 11.93 ± 8.27 (12) | 16.65 ± 12.53 (11) | 9.93 ± 1.31 (10) | 1.29 ± 0.20<sup>a</sup> (12) | 1.09 ± 0.25<sup>b</sup> (11) | 1.07 ± 0.23<sup>b</sup> (10) |
| Kidney | 6.77 ± 1.16<sup>a</sup> (12) | 11.33 ± 4.18<sup>b</sup> (13) | 6.87 ± 1.17<sup>a</sup> (8) | 1.13 ± 0.16<sup>a</sup> (10) | 1.38 ± 0.26<sup>b</sup> (13) | 1.20 ± 0.19<sup>a</sup> (8) |
| Muscle | ND (9) | ND (8) | ND (8) | 0.87 ± 0.41 (9) | 0.84 ± 0.31 (8) | 0.61 ± 0.17 (8) |
| BAT | 25.87 ± 16.37 (8) | 25.63 ± 10.00 (8) | 28.80 ± 7.52 (7) | 1.88 ± 0.31 (8) | 1.70 ± 0.42 (8) | 1.69 ± 0.31 (7) |
| WAT | ND (5) | 4.10 ± 1.05 (3) | 4.89 ± 0.83 (3) | 0.50 ± 0.34<sup>ab</sup> (5) | 0.33 ± 0.01<sup>a</sup> (3) | 0.44 ± 0.01<sup>b</sup> (3) |

Note: Tissues of mice at the age of 11 weeks are used for gas chromatography, and 19:0 is an internal standard. ND, Not detected. Each value represents the mean ± SD, and the numbers in brackets indicate the size of the tested samples. For all fatty acid contents, see Supplementary Table 4. Different letters in superscript indicate a level of statistical significance of $p < 0.05$ .

The contents of other FAs, such as myristic (14:0), palmitic (16:0), palmitoleic (16:1n-7), stearic (18:0), oleic (18:1n-9), cis-vaccenic (18:1n-7), or arachidonic (20:4n-6) acids, were changed by varying degrees in pai heart, kidney, liver or WAT tissues (Supplementary Table 4). In contrast, no evident changes in FA compositions were observed in pai muscle and BAT tissue, suggesting the content changes of some FAs are genotype-or tissue-specific. Simultaneously, the amounts of total FAs were decreased in the pai/pai livers (9.19 ± 1.55 in wt versus 8.19 ± 1.78 mg/g in pai/pai; p < 0.05) and contrarily increased in both pai kidneys (9.59 ± 0.99 in wt versus 12.97 ± 2.21 in pai/wt versus 10.66 ± 1.06 in pai/pai; p < 0.05; Supplementary Table 4), suggesting fat loss in livers but fat accumulation in kidneys.

### 3.3. Weaning and growth analysis

Considering the potential impact of maternal t10c12-CLA from the placenta and/or milk, pups delivered by wt or pai/wt mothers were measured separately. In the group of wt mothers (Fig. 2A), the pai/wt pups showed no bodyweight difference from the wt littermates during the post-weaning development. In the group of pai/wt mothers (Fig. 2B), the pai/wt or pai/pai pups also showed no bodyweight difference from the wt littermates during the post-weaning development. However, the pai/wt genotypes were overweight (p < 0.05) compared to pai/pai mice from 10 weeks onwards.

**Figure 2.**
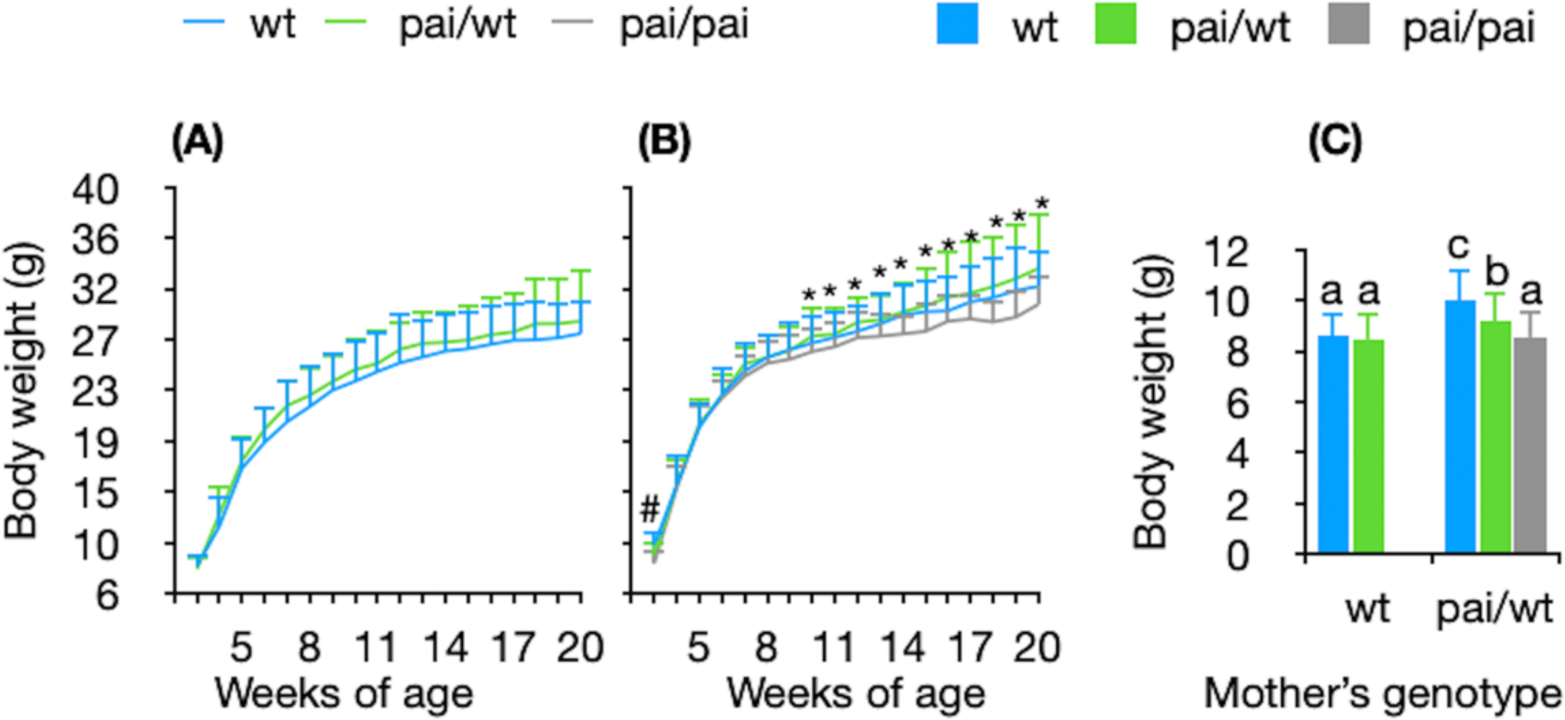
Growth curves of different genotype pups from wt (A) or pai/wt (B) mother and comparison of the body weight (C) of pups at the age of 3 weeks. Each group contained 15-20 (A, B) or 30-85 (C) pups. Bars represent the mean ± SD. # and different letters indicate p < 0.05 among three genotypes; * indicates p < 0.05 between pai/pai and pai/wt genotypes.

Interestingly, within the group of pai/wt mothers, the weaning weights of three genotypes were significantly different (pai/pai < pai/wt < wt, respectively; p < 0.05. Fig. 2 B-C), suggesting the endogenous t10c12-CLA produced by foetus/pups themselves would result in the reduced bodyweight during embryonic/weaning development. On the other hand, the weaning weights of wt or pai/wt pups from pai/wt mothers were respectively overweight than that from wt mother (p < 0.05; Fig. 2C), suggesting that the maternal t10c12-CLA might result in the increased bodyweight of the embryos/pups.

### 3.4. Energy metabolism and activity studies

Energy metabolism and activity parameters were determined under the consideration of body weight in the 11-week-old pai/pai mice and wt littermates over 72 hrs in metabolic cages (Fig. 3). Compared to the wt littermates, pai/pai mice showed no different (p > 0.05) changes in food intake, respiratory exchange ratio, marginal activities, and speed in both light and dark phases; as well as body weight gain and total distance travelled during the 72 hrs period.

**Figure 3.**
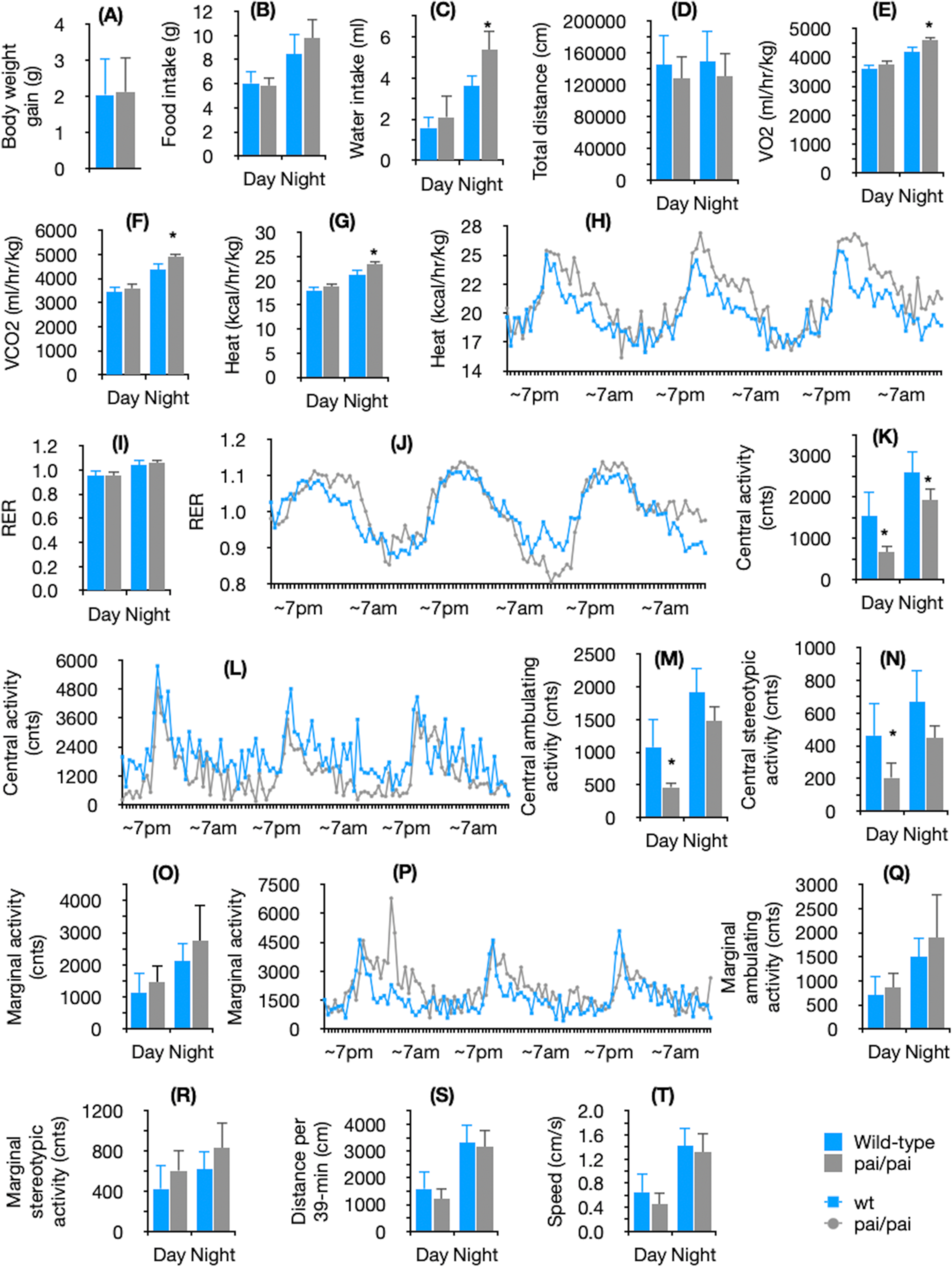
Pai mice have abnormal energy metabolism and activities. Comparison of body weight gain during 72 h period (A), food intake (B), water intake (C), total distance travelled during 72 h period (D), O_2_ consumption (E), CO_2_ production (F), heat release and its time course (G-H), respiratory exchange ratio (REF) and RER time course (I-J), locomotor activities in the centre of the cage and its time course (K-L), central ambulating and stereotypic activities (M-N), locomotor activities in the margin or corners of the cage and its time course (O-P), marginal ambulating and stereotypic activities (Q-R), distance travelled during a 39-min sampling duration (S), and speed (T) among wt (n=5) and pai/pai (n=5) mice at the age of 11 weeks. Lights turn on and off at 7:00 am and 7:00 pm, respectively. The data are normalised to lean body weight. Bars represent the mean ± SD. * indicate p < 0.05 within the same phase.

However, pai/pai mice had significantly (p < 0.05) fewer central activities in both light (56%) and dark (26%) phases, but they consumed 32% and 48% more water, 4.1% and 9.9% more O_2_, and produced 4.3% and 12% more CO_2_, and 4.2% and 10.4% more heat during the light (p > 0.05) and dark (p < 0.05) periods, respectively (Fig. 3). That was, pai/pai mice reduced their activities, but consumed more oxygen (energy) to release more heat for maintenance.

### 3.5. Metabolic complications

To match energy demand and expenditure, the circulating hormonal signals were examined under the non-fasted state. The concentrations of serum insulin, leptin, ghrelin, and circulating glucose (Table 2), as well as the length of the small intestine (Supplementary Table 5), showed no differences (p > 0.05) among the three genotypes. It suggests that the t10c12-CLA does not play a role in the energy intake, consistent with the observation in the metabolic cage.

**Table 2.**
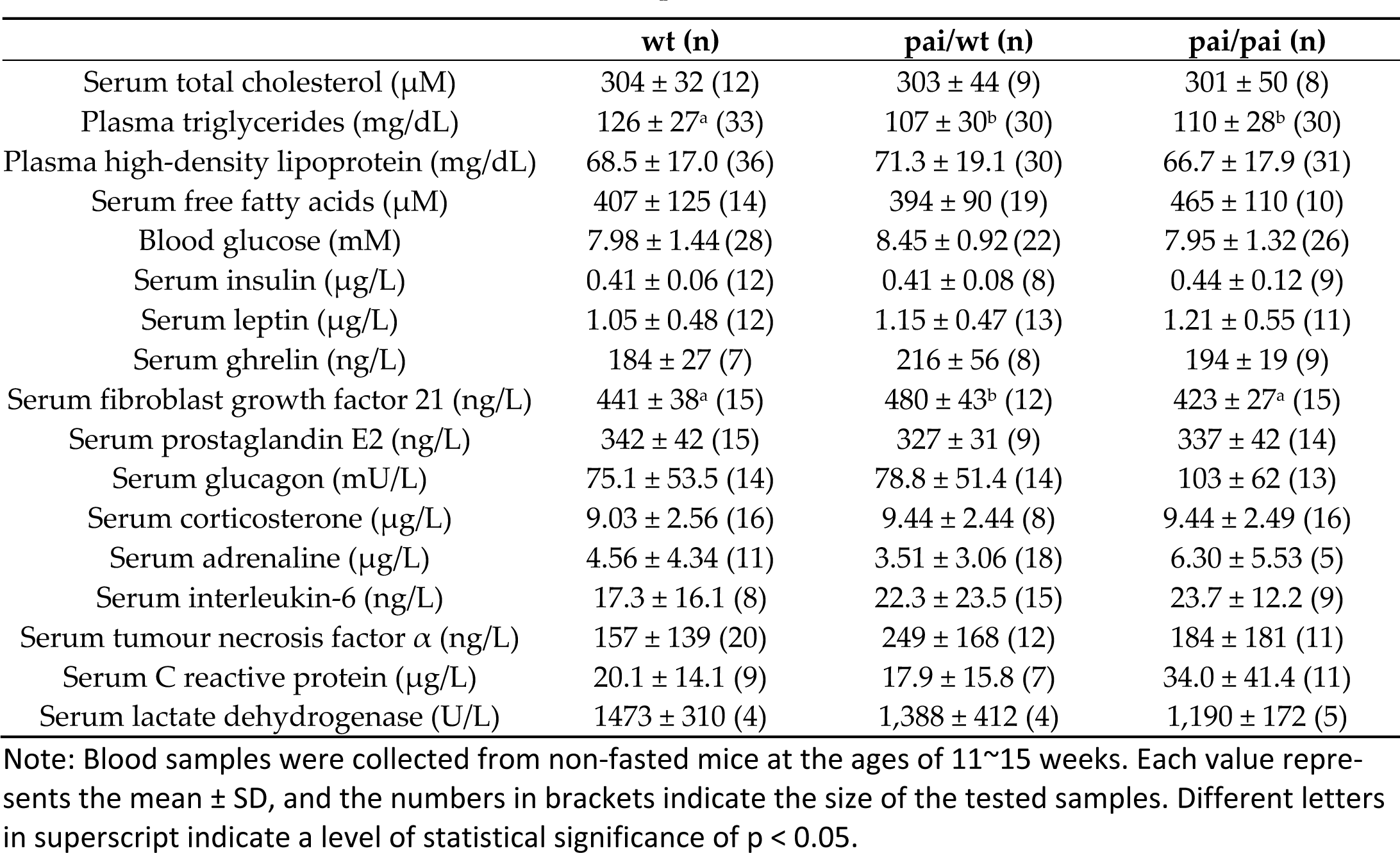
Hormones/factors in blood in wt and pai mice.

|  | wt (n) | pai/wt (n) | pai/pai (n) |
| --- | --- | --- | --- |
| Serum total cholesterol (μM) | 304 ± 32 (12) | 303 ± 44 (9) | 301 ± 50 (8) |
| Plasma triglycerides (mg/dL) | 126 ± 27<sup>a</sup> (33) | 107 ± 30<sup>b</sup> (30) | 110 ± 28<sup>b</sup> (30) |
| Plasma high-density lipoprotein (mg/dL) | 68.5 ± 17.0 (36) | 71.3 ± 19.1 (30) | 66.7 ± 17.9 (31) |
| Serum free fatty acids (μM) | 407 ± 125 (14) | 394 ± 90 (19) | 465 ± 110 (10) |
| Blood glucose (mM) | 7.98 ± 1.44 (28) | 8.45 ± 0.92 (22) | 7.95 ± 1.32 (26) |
| Serum insulin (μg/L) | 0.41 ± 0.06 (12) | 0.41 ± 0.08 (8) | 0.44 ± 0.12 (9) |
| Serum leptin (μg/L) | 1.05 ± 0.48 (12) | 1.15 ± 0.47 (13) | 1.21 ± 0.55 (11) |
| Serum ghrelin (ng/L) | 184 ± 27 (7) | 216 ± 56 (8) | 194 ± 19 (9) |
| Serum fibroblast growth factor 21 (ng/L) | 441 ± 38<sup>a</sup> (15) | 480 ± 43<sup>b</sup> (12) | 423 ± 27<sup>a</sup> (15) |
| Serum prostaglandin E2 (ng/L) | 342 ± 42 (15) | 327 ± 31 (9) | 337 ± 42 (14) |
| Serum glucagon (mU/L) | 75.1 ± 53.5 (14) | 78.8 ± 51.4 (14) | 103 ± 62 (13) |
| Serum corticosterone (μg/L) | 9.03 ± 2.56 (16) | 9.44 ± 2.44 (8) | 9.44 ± 2.49 (16) |
| Serum adrenaline (μg/L) | 4.56 ± 4.34 (11) | 3.51 ± 3.06 (18) | 6.30 ± 5.53 (5) |
| Serum interleukin-6 (ng/L) | 17.3 ± 16.1 (8) | 22.3 ± 23.5 (15) | 23.7 ± 12.2 (9) |
| Serum tumour necrosis factor $\alpha$ (ng/L) | 157 ± 139 (20) | 249 ± 168 (12) | 184 ± 181 (11) |
| Serum C reactive protein (μg/L) | 20.1 ± 14.1 (9) | 17.9 ± 15.8 (7) | 34.0 ± 41.4 (11) |
| Serum lactate dehydrogenase (U/L) | 1473 ± 310 (4) | 1,388 ± 412 (4) | 1,190 ± 172 (5) |

Note: Blood samples were collected from non-fasted mice at the ages of 11~15 weeks. Each value represents the mean ± SD, and the numbers in brackets indicate the size of the tested samples. Different letters in superscript indicate a level of statistical significance of $p < 0.05$ .

Either glucose or insulin tolerance tests showed no difference in the circulating glucose concentrations or the area under the curve between wt and two pai genotypes (Fig. 4 A-B). However, the blood glucose levels before glucose injection were remarkably (p < 0.05; Fig. 4 A) increased in pai/wt mice and maintained typically when fasted 24-h duration (Fig. 4 B). It suggests that the t10c12-CLA might ameliorate the glucose sensitivity in pai/wt mice at the beginning of starvation, which may be associated with the elevated FGF21 in pai/wt mice (p < 0.05; Table 2) for FGF21 plays an active role in insulin-sensitising and energy expenditure [23].

**Figure 4.**
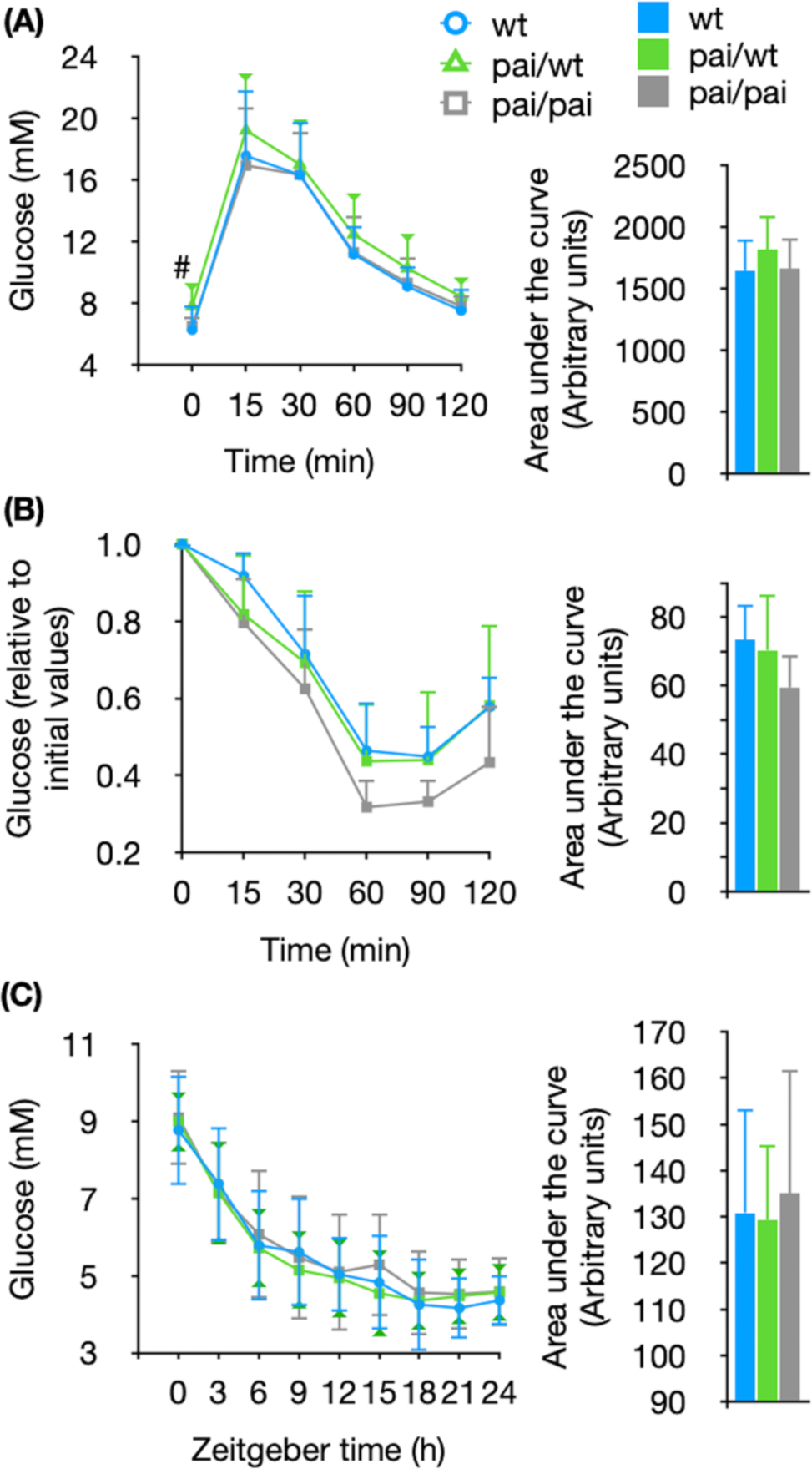
Glucose (A) and insulin (B) tolerance tests, and dynamic glucose levels (C) in wt and pai mice. (A) Comparison of absolute glucose levels and the area under curve among wt (n = 13), pai/wt (n = 19), and pai/pai (n = 10) mice. (B) Comparison of blood glucose levels relative to initial values and the area under the curve among wt (n = 4), pai/wt (n = 8), and pai/pai (n =4) mice. (C) Dynamic blood glucose changes and the area under the curve in wt (n = 8), pai/wt (n = 11), and pai/pai (n = 8) fasting mice starved from Zeitgeber time 0 to 24. Zeitgeber times 0 and 12 are defined as lights-on and-off times, respectively. Bars represent the mean ± SD. # indicates p < 0.05 between pai/wt with wt or pai/pai mice.

In addition, there were no differences in the circulating TC, FFAs, and HDL among the three genotypes, but the serum levels of TGs were reversely decreased in pai/wt (p = 0.011) or pai/pai mice (p = 0.021; Table 2) compared to wt mice.

Furthermore, the serum PGE2, glucagon, corticosterone, adrenaline, lactate dehydrogenase, TNFα, IL-6, and CRP in pai/wt or pai/pai mice were respectively kept at normal levels compared with wt mice (p > 0.05; Table 2), suggesting no pathological symptoms such as cellular toxicity of expressing the bacterial protein, response to illness-related stimuli, and inflammation in pai mice. It also indicates that the phenotype profiling of pai mice results from the t10c12-CLAs’ impact.

### 3.6. WAT reduction

Mice at 11 weeks were used to determine the WAT features by dissection, magnetic resonance (MR) imaging, histological, and RNA analyses. The results were genotype-specific and described as follows. Compared to wt mice, in pai/wt mice, no evident changes in organs/tissues, adipocyte volume or cross-sectional areas per adipocyte (Fig. 5 B-D) were observed. On the contrary, in pai/pai mice, organs such as livers, spleens, and kidneys were significantly enlarged (p < 0.05; Supplementary Table 5). Simultaneously, the white fat was lost (Fig. 5 A-B), consistent with the reduced adipocyte volume or cross-sectional areas per adipocyte (Fig. 5 C-D), as well as the lowered RNA levels of leptin, Cebpβ (Full name seen in Table 3) that plays a role in the early stages of adipogenesis, macrophage markers F4/80, CD68, and CD11c in WAT (Table 3). The results suggest that the t10c12-CLA can result in fat loss in pai/pai mice.

**Figure 5.**
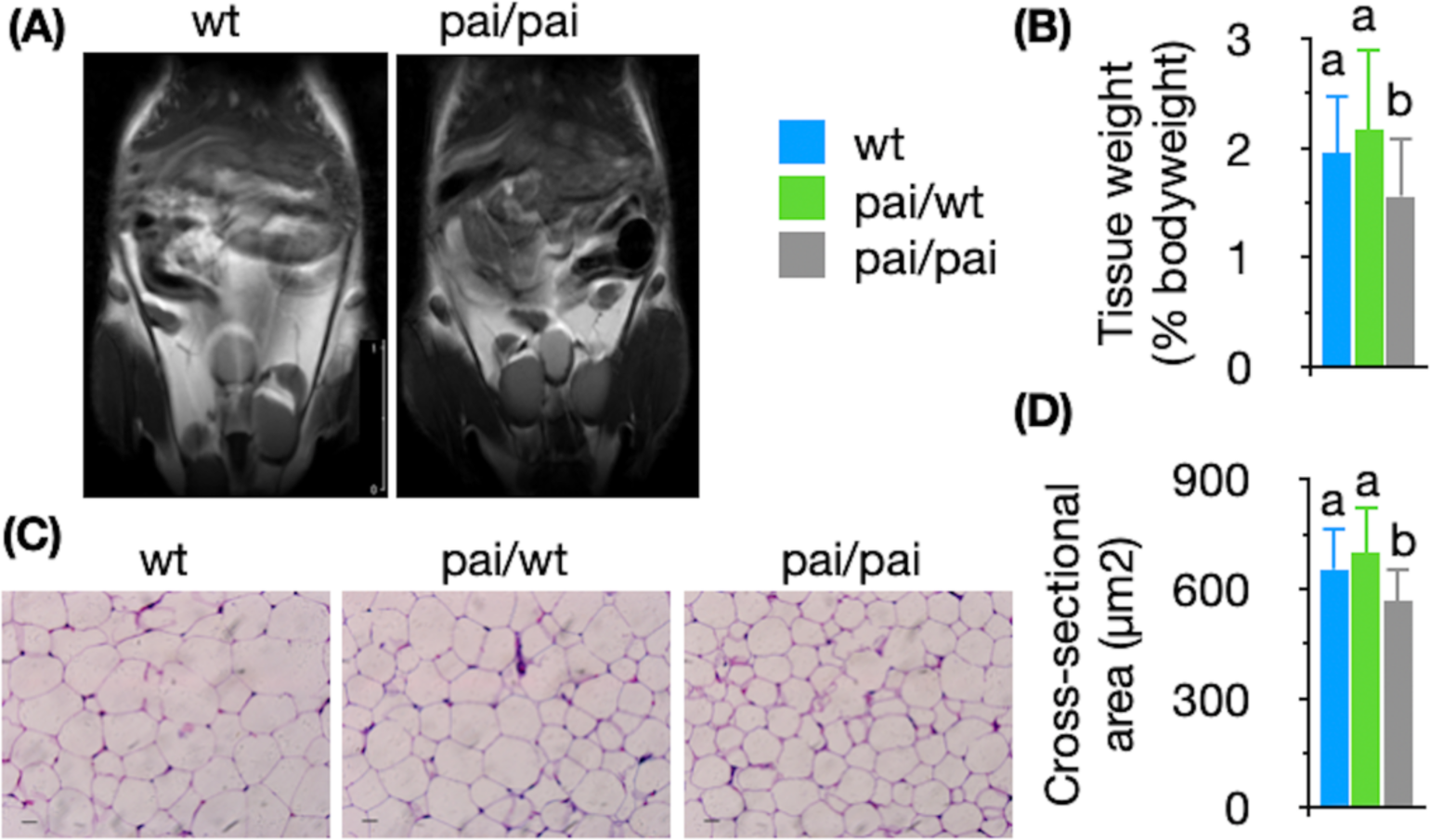
Magnetic resonance imaging and histological analysis of white adipose tissues. (A) Coronal sections were obtained from the whole-body midsection for five wt or pai/pai mice at 11 weeks in each group during magnetic resonance imaging (bar = 1 cm). The wt mice show adipose tissue in the subcutaneous and intra-abdominal regions as areas of increased signal intensity on images. The pai/pai mice show less fat. (B) The WAT weight and (C) Imaging by hematoxylin-eosin staining (bar = 10 µm), as well as (D) analysis of cross-sectional area per adipocyte, show that the volumes of white adipocytes are reduced in pai/pai mice. Bars represent the mean ± SD. Different letters indicate p < 0.05.

**Table 3.**
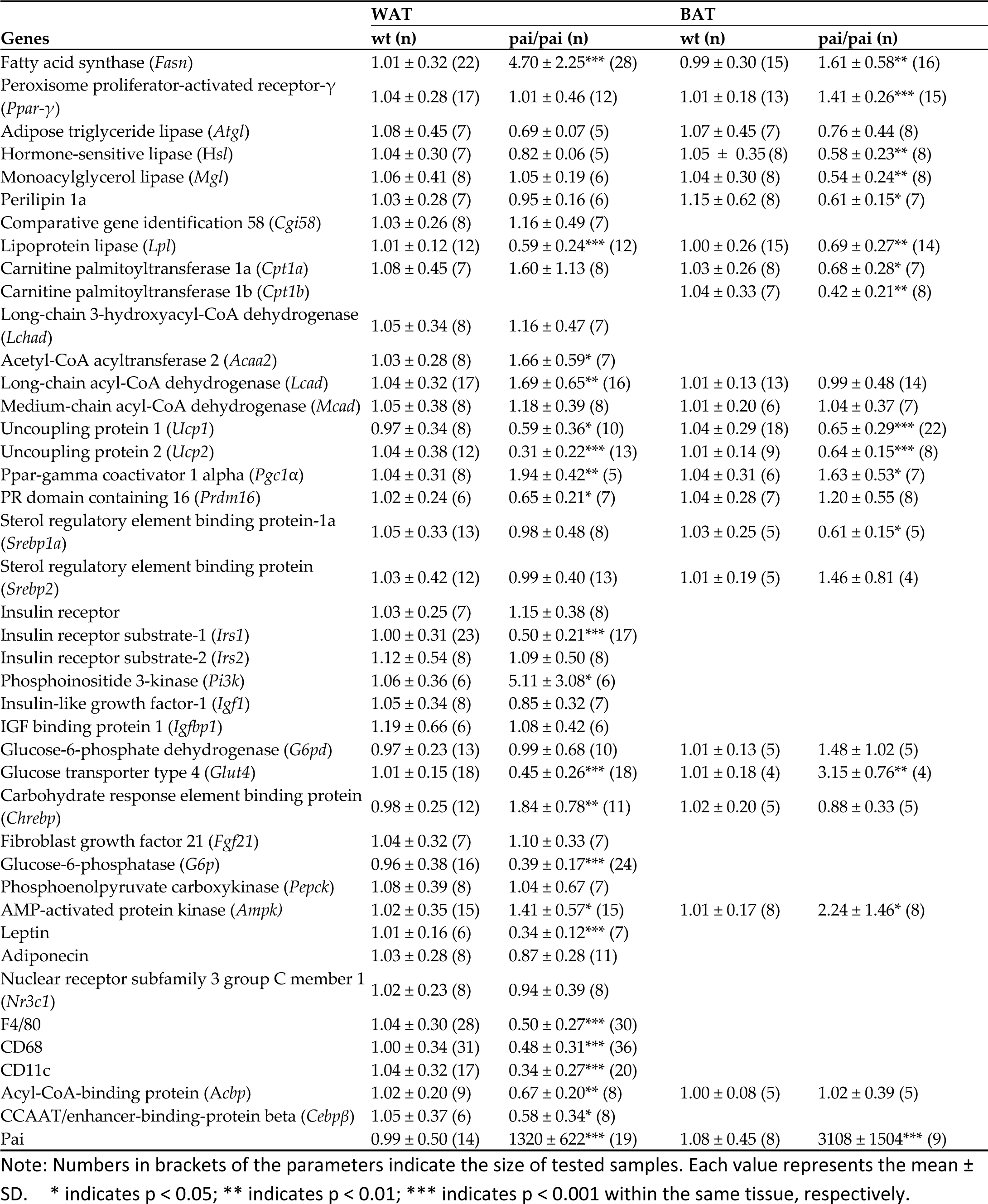
Relative expression of mRNAs in the white (WAT) and brown (BAT) adipose tissues.

In pai/pai white adipocytes, significant (p < 0.05) changes in RNA levels appeared in 50% (20/40) tested genes of which seven were up-regulated and 13 were down-regulated (Table 3). The energy regulator Ampk was up-regulated, suggesting the inhibition of lipid synthesis, which is consistent with the down-regulated lipoprotein lipase that supports the uptake of FAs from very low-density lipoproteins in the bloodstream, regular expression of Srebp1a and Srebp2 involved in the sterol pathway and transcriptional factor Ppar-γ, while the fatty acid synthase was over-expressed. Genes involved in lipolyses, such as Atgl, hormone-sensitive lipase, Mgl, Perilipin 1a, and Ggi58 kept normal RNA levels. Normal RNA levels of genes involved in beta-oxidation, including Cpt1a, Lchad, and Mcad and the reduced RNA levels of the acyl-CoA-binding protein that act as an intracellular carrier of acyl-CoA esters were detected in pai/pai WAT, suggesting the weakening process of beta-oxidation in mitochondria. On the contrary, the RNA levels of Acaa2 and Lcad were increased, suggesting the up-regulation of beta-oxidation of very long-chain FAs in peroxisomes.

In pai/pai adipocytes, although Pgc1α, which activates mitochondrial biogenesis genes through interaction with Ppar-γ during energy stress, were up-regulated. Simultaneously, Prdm16, Ucp1, and Ucp2 were down-regulated, suggesting the declined WAT browning.

Consistent with the normal levels of the circulating insulin or glucocorticoids, their receptor genes Insr and Nr3c1, also expressed commonly on the pai/pai adipocyte surface; interestingly, the Irs1 was down-regulated while its downstream Pi3K gene was up-regulated. The RNA levels of other genes involved in insulin/IGF-1 signalling, such as Irs2, Igf1, and Igfbp1 showed no changes compared to the wt samples. Meanwhile, there were reduced expression of Chrebp and glucose transporter Glut4 and up-regulated glucose-6-phosphatase involved in gluconeogenesis, suggesting the energy mobilisation for glucose homeostasis. These results indicate that the t10c12-CLA has induced the broadly positive or negative feedback regulation of lipid and glucose metabolism in WAT.

### 3.7. BAT thermoregulation

Parameters of BAT were measured to investigate the molecular mechanism of more heat production in pai mice. MR imaging and dissection analysis revealed that the weight (% bodyweight) of interscapular BAT was increased from 0.39 ± 0.07 in wt to 0.50 ± 0.18 in pai/wt (p = 0.059) and 0.54 ± 0.18 in pai/pai mice (p < 0.05; Fig. 6 A-C). HE staining showed that there were more small-sized and irregular lipid drops within the pai/wt or pai/pai adipocytes, quite different from the wt adipocytes containing relatively uniform and medium-sized lipid drops (Fig. 6 D); simultaneously, the size of cross-sectional area per brown adipocyte was enlarged in both pai genotypes (p < 0.05; Fig. 6 E). It suggests that t10c12-CLA-induced BAT mass increased in pai mice.

**Figure 6.**
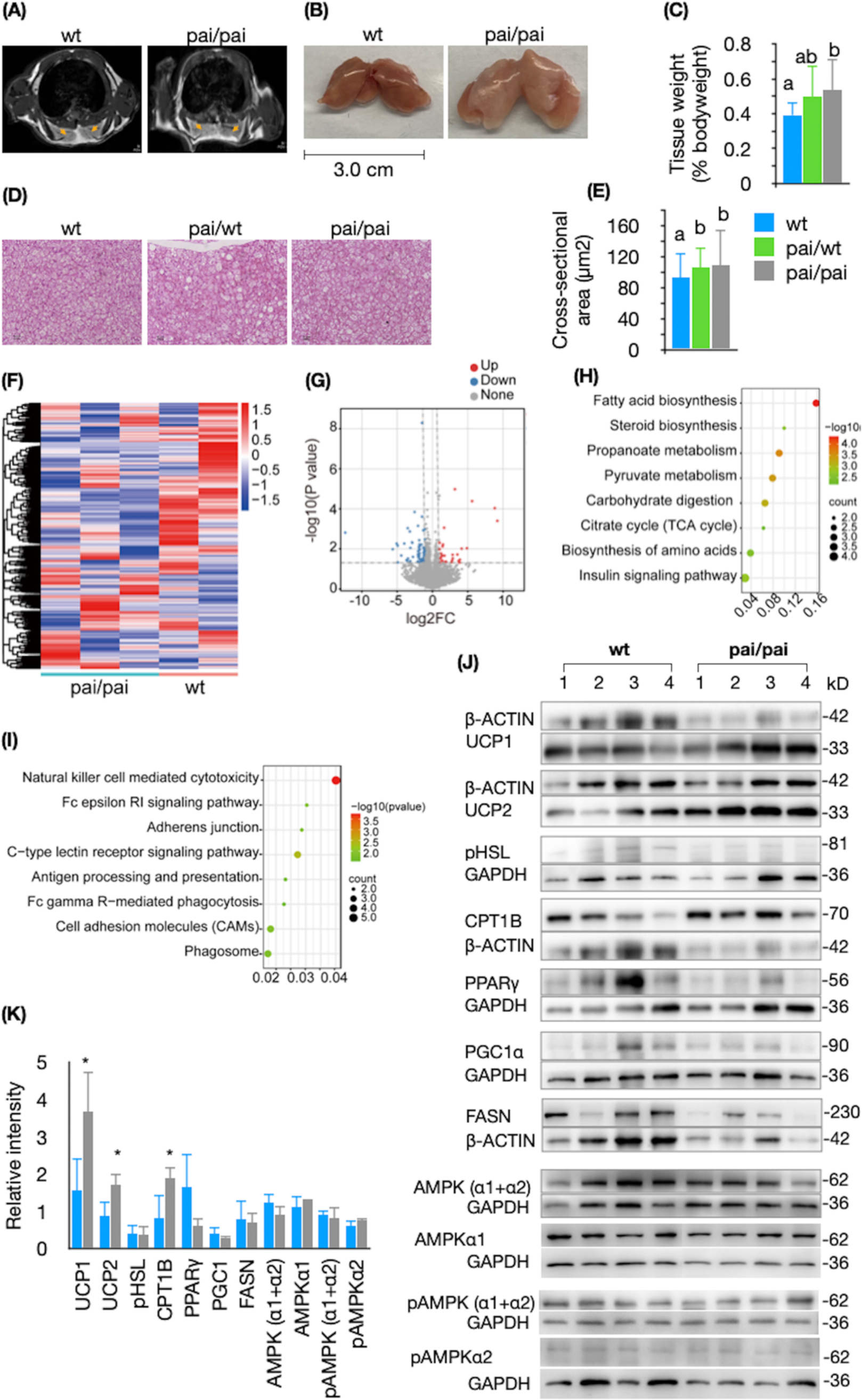
Aspects of brown adipose tissues. Axial sections of the interscapular BAT show more prominent areas of grey signal intensity on magnetic resonance images (Brown arrows) in five pai/pai mice than in five wt mice at 11 weeks (A). The BAT mass increases in pai/pai mice (B, C). Imaging by hematoxylin-eosin staining show more small-sized and irregular lipid drops in the pai/wt or pai/pai adipocytes and relatively uniform and medium-sized lipid drops in the wt adipocytes (D, bar = 10 µm) and analysis of cross-sectional area per cell show that the volumes of brown adipocytes are increased in pai mice (E). RNA-Seq analysis shows transcriptional changes of differentially expressed genes in BAT from two wt and three pai/pai mice using heatmap (F) and the up-or down-regulated genes in pai/pai BAT using volcano plots (G). Maps of KEGG enrichment analysis show the up-(H) and down-regulated (I) metabolic pathways in pai/pai BAT. Western blot images (J) and relative intensities (K) analyses of some critical proteins in the BAT from wt and pai/pai mice. pAMPK ⍺1 or ⍺2 indicate phosphorylation takes place at T183 (AMPK α1, phosphorylated at threonine 183) and T172 (AMPK α2, phosphorylated threonine 172), respectively. CPT1B, carnitine palmitoyltransferase 1b; FASN, fatty acid synthase; PGC1A, pparγ coactivator 1α; pHSL, phosphorylated hormone-sensitive lipase; PPARγ, peroxisome proliferator-activated receptor-γ; UCP, uncoupling protein. Bars represent the mean ± SD. Different letters in superscript or * indicate p < 0.05.

RNA sequencing provided a snapshot of the transcriptional dynamics between the wt and pai/pai BAT. The heatmap (Fig. 6 F) and a volcano plot (Fig. 6 G) revealed that the global transcriptional profile of pai/pai BAT differed from that of wt BAT. KEGG pathway enrichment analysis showed that the up-regulated pathways in pai/pai BAT included fatty acid biosynthesis, pyruvate metabolism, carbohydrate digestion and absorption, Insulin signalling pathway, steroid biosynthesis, biosynthesis of amino acids, citrate cycle, and propanoate metabolism (Fig. 6 H). In contrast, the down-regulated pathway included NK cell-mediated cytotoxicity, C-type lectin receptor signalling pathway, cell adhesion molecules, phagosome, tuberculosis, Fc epsilon RI signalling pathway, adherens junction, antigen processing and presentation, and Fc gamma R-mediated phagocytosis signalling pathways (Fig. 6 I).

Based on the prediction of transcriptome analysis and the conventional physiological factors that can impact BAT’s activity, 24 genes were chosen for real-time PCR analysis and fourteen (56%) of them altered their transcriptional patterns in pai/pai brown adipocytes. Among them, five upregulated genes were Ampk, Fasn, Glut4, Ppar-r, and Pgc1α and nine down-regulated genes were Srebp1a, Lpl, Hsl, Mgl, Perilipin1a, Cpt1a, Cpt1b, Ucp1, and Ucp2 (Table 3). In addition, western blot analysis revealed that there was no difference in protein levels of AMPK, pAMPK on Thr172 or Thr183, PGC1A, FASN, PPAR-γ, and pHSL in the samples of BAT from the pai/pai and wt mice (Fig. 6 J-K), suggesting there was no AMPK activation rewired metabolism to decrease anabolic processes and increase catabolism. However, the elevation of CPT1B, UCP1 and UCP2 proteins suggests the increased lipid beta-oxidation and BAT thermogenesis in pai/pai mice. The inconsistency between some RNA and protein levels suggests the possibility of positive or negative feedback regulation in pai mice.

### 3.8. Hepatic features

Compared to wt mice, the hepatic analysis also showed varying differences between pai/wt and pai/pai livers. Besides liver hypertrophy (p < 0.05; Fig. 7 A) and the amount decrease of total FAs (p < 0.05; Supplementary Table 4) in pai/pai livers, the hepatic TC or TGs levels had not changed (p > 0.05; Fig. 7 B-C), and no steatosis was observed by histological staining of liver slices from pai/wt and pai/pai mice, while the estimation of cross-sectional area per cell showed swollen hepatocytes (Fig. 7 D-G), consistent with liver hypertrophy in both pai genotypes. In addition, the mRNA levels of 58% (28/48) critical genes were significantly modified in pai/pai livers, including 11 down-regulated and 17 up-regulated genes (p < 0.05; Table 4), suggesting that the t10c12-CLA might influence the liver metabolism in mice.

**Figure 7.**
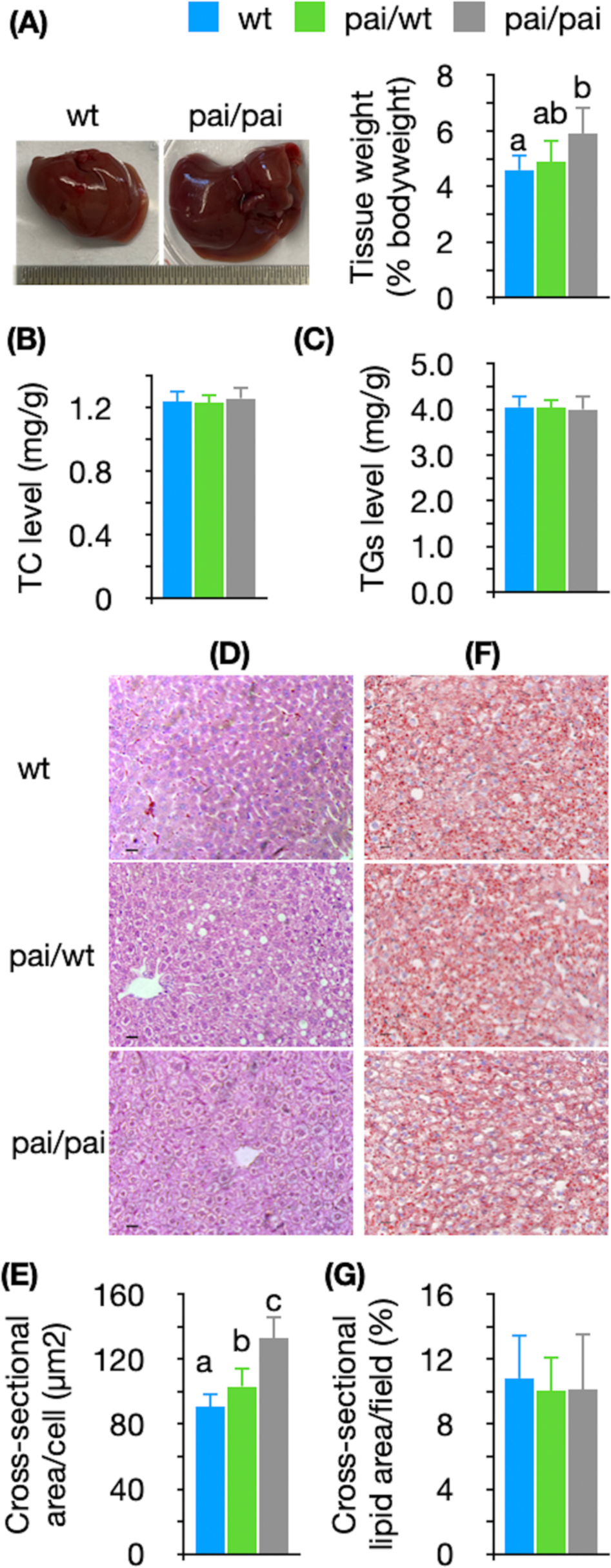
Lipid and histological analysis of livers. Comparison of weight (A), total cholesterol (B), and triglycerides (C) levels in wt and pai livers. Imaging of hematoxylin-eosin staining shows the abnormal morphology and oedema of pai hepatocytes. Analysis of cross-sectional area per cell indicates that the cellular volumes gradually enlarged in pai/wt and pai/pai hepatocytes, respectively (D-E, Bar = 10 µm). Oil red staining and analysis of the cross-sectional area of lipids per field show no lipid accumulation in either pai liver (F∼G). Bars represent the mean ± SD; different letters indicate p < 0.05.

**Table 4.** Relative expression of mRNAs in the livers.

| Genes | wt (n) | pai/pai (n) |
| --- | --- | --- |
| *Fatty acid and TGs synthesis* |  |  |
| Fatty acid synthase (*Fasn*) | 1.03 ± 0.26 (14) | 1.72 ± 0.56*** (13) |
| Stearoyl-CoA desaturase 1 (*Scd1*) | 1.06 ± 0.41 (7) | 1.19 ± 0.50 (5) |
| Fatty acid desaturase 1 (*Fads1*) | 1.01 ± 0.21 (3) | 1.08 ± 0.18 (3) |
| Fatty acid desaturase 2 (*Fads2*) | 1.04 ± 0.33 (3) | 0.92 ± 0.26 (3) |
| Peroxisome proliferator-activated receptor-γ (*Ppar-γ*) | 1.04 ± 0.35 (28) | 0.65 ± 0.31*** (23) |
| Glycerol-3-phosphate acyltransferase 1 (*Gpat1*) | 1.05 ± 0.35 (7) | 1.04 ± 0.37 (8) |
| 1-acylglycerol-3-phosphate O-acyltransferase 2 (*Agpat2*) | 1.02 ± 0.18 (15) | 1.11 ± 0.25 (14) |
| Diacylglycerol acyltransferase-1 (*Dgat1*) | 1.04 ± 0.28 (17) | 1.37 ± 0.39* (15) |
| Diacylglycerol acyltransferase-2 (*Dgat2*) | 1.03 ± 0.26 (9) | 1.05 ± 0.32 (9) |
| CD36 | 1.07 ± 0.38 (8) | 0.58 ± 0.32* (8) |
| ***Lipolysis and fatty acid oxidation*** |  |  |
| Hepatic triglyceride lipase (*Htgl*) | 1.06 ± 0.37 (9) | 1.25 ± 0.28 (9) |
| Adipose triglyceride lipase (*Atgl*) | 1.04 ± 0.33 (9) | 1.03 ± 0.36 (9) |
| Hormone-sensitive lipase (*Hsl*) | 1.07 ± 0.38 (9) | 1.08 ± 0.26 (9) |
| Comparative gene identification 58 (*Cgi58*) | 1.03 ± 0.27 (9) | 1.40 ± 0.38* (9) |
| Lipoprotein lipase (*Lpl*) | 0.99 ± 0.16 (11) | 1.37 ± 0.15*** (6) |
| Carnitine palmitoyltransferase Ia (*Cpt1a*) | 1.05 ± 0.37 (9) | 1.01 ± 0.35 (9) |
| Long-chain acyl-CoA dehydrogenase (*Lcad*) | 1.01 ± 0.18 (14) | 1.11 ± 0.49 (14) |
| Medium-chain acyl-CoA dehydrogenase (*Mcad*) | 1.06 ± 0.38 (15) | 0.95 ± 0.36 (13) |
| Uncoupling protein 1 (*Ucp1*) | 1.00 ± 0.31 (20) | 0.59 ± 0.36** (13) |
| Uncoupling protein 2 (*Ucp2*) | 1.04 ± 0.28 (16) | 0.82 ± 0.28* (14) |
| ***Sterol pathway*** |  |  |
| Liver X receptor α (*Lxrα*) | 1.01 ± 0.16 (9) | 1.15 ± 0.40 (9) |
| Liver X receptor β (*Lxrβ*) | 1.02 ± 0.23 (9) | 1.22 ± 0.47 (9) |
| Sterol Regulatory Element Binding Protein-1a (*Srebp1a*) | 1.07 ± 0.39 (7) | 1.54 ± 0.28* (7) |
| Sterol Regulatory Element Binding Protein-1c (*Srebp1c*) | 1.12 ± 0.69 (4) | 1.29 ± 0.44 (5) |
| Sterol Regulatory Element Binding Protein-2 (*Srebp2*) | 1.00 ± 0.30 (16) | 1.40 ± 0.71* (18) |
| HMG-CoA reductase (*Hmgcr*) | 1.03 ± 0.27 (6) | 0.65 ± 0.28* (7) |
| ***Insulin and IGF-1 signaling*** |  |  |
| Insulin receptor (*Insr*) | 0.99 ± 0.24 (16) | 1.40 ± 0.34*** (17) |
| Insulin receptor substrate-1 (*Irs1*) | 1.03 ± 0.28 (29) | 1.19 ± 0.68 (25) |
| Insulin receptor substrate-2 (*Irs2*) | 1.02 ± 0.34 (15) | 1.55 ± 0.52** (16) |
| Insulin-like growth factor-1 (*Igf1*) | 1.01 ± 0.23 (21) | 1.40 ± 0.55** (22) |
| IGF binding protein 1 (*Igfbp1*) | 1.01 ± 0.23 (15) | 0.53 ± 0.26*** (18) |
| Forkhead box protein a2 (*Foxa2*) | 1.03 ± 0.31 (27) | 1.75 ± 0.77*** (30) |
| Igf1 receptor (*Igf1r*) | 1.07 ± 0.37 (5) | 1.63 ± 0.28* (4) |
| ***NADPH-producing*** |  |  |
| Malic enzyme | 1.02 ± 0.23 (8) | 1.89 ± 0.93* (7) |
| Glucose-6-phosphate dehydrogenase (*G6pd*) | 0.98 ± 0.19 (14) | 2.34 ± 0.63*** (17) |
| 6-Phosphogluconate dehydrogenase (*Pgd*) | 1.24 ± 0.93 (8) | 1.29 ± 0.80 (9) |
| ***Glucose metabolism and gluconeogenesis*** |  |  |
| Glucose transporter type 4 (*Glut4*) | 0.99 ± 0.23 (20) | 0.59 ± 0.23*** (15) |
| Glucokinase | 1.02 ± 0.23 (12) | 0.66 ± 0.28** (13) |
| Carbohydrate response element binding protein (*Chrebp*) | 1.00 ± 0.28 (13) | 1.98 ± 0.60*** (11) |
| Fibroblast growth factor 21 (*Fgf21*) | 1.32 ± 1.07 (9) | 1.34 ± 0.91 (8) |
| Glucose-6-phosphatase (*G6p*) | 1.02 ± 0.19 (14) | 2.34 ± 0.63*** (25) |
| Phosphoenolpyruvate carboxykinase (*Pepck*) | 1.06 ± 0.35 (13) | 0.69 ± 0.31** (14) |
| *Others* |  |  |
| AMP-activated protein kinase (*Ampk*) | 1.06 ± 0.37 (9) | 1.07 ± 0.35 (9) |
| Nuclear receptor subfamily 3 group C member 1 (*Nr3c1*) | 1.03 ± 0.31 (3) | 1.26 ± 0.36 (3) |
| F4/80 | 1.02 ± 0.20 (14) | 1.45 ± 0.53* (13) |
| CD68 | 1.05 ± 0.34 (10) | 0.76 ± 0.22* (10) |
| CD11c | 1.05 ± 0.31 (19) | 1.93 ± 0.89** (17) |
| Forkhead box protein c2 (*Foxc2*) | 1.01 ± 0.17 (8) | 1.59 ± 1.02 (10) |
| Pai | 1.17 ± 0.69 (7) | 10830 ± 8060* (7) |

Numbers in brackets of the parameters indicate the size of tested samples. Each value represents the mean ± SD. \* indicates $p < 0.05$ ; \*\* indicates $p < 0.01$ ; \*\*\* indicates $p < 0.001$ , respectively.

Although the hepatic TC and TGs levels were normal in pai/pai mice, their hepatic Srebp1a and Srebp2 were up-regulated, and the HMG-CoA reductase (Hmgcr) was down-regulated. Some genes involved in FAs/TGs synthesis had also altered their transcription levels. For instance, the RNA levels of Fasn, Dgat1 which convert diacylglycerol into TGs, and Lpl which support FAs uptake, were increased; and the RNA levels of Ppar-γ and CD36, which transport long-chain FAs into the cells, were decreased. Simultaneously, a series of crucial enzymes involved in lipolysis and beta-oxidation kept standard transcription except for the up-regulated Cgi58. In addition, the Ucp 1 and 2 engaged in thermogenesis were down-regulated; the malic enzyme and G6pd, which both produce NADPH, were significantly up-regulated; the inflammatory factors F4/80 and CD11c were up-regulated, but CD68 was down-regulated (Table 4).

For genes involved in insulin/IGF1 signalling, except for reduced Igfbp1 and commonly expressed Irs1, the other five essential genes, including Insr, Irs2, Igf1, Igf1 receptor, and pioneering transcription factor Foxa2 were elevated altogether. For genes involved in glucose metabolism and gluconeogenesis, the RNA levels of Glut4, glucokinase, and Pepck were reduced, whereas the RNA levels of G6p and Chrebp were elevated. These transcriptional changes suggest that t10c12-CLA easily influences hepatic glucose metabolism and insulin sensitivity.

### 3.9. Hypothalamic gene analysis

To determine whether gene expression has been affected by t10c12-CLA in the hypothalamus, which contains highly conserved neural circuitry controlling energy metabolism, fluid and electrolyte balance, thermoregulation, responses to stressors, wake-sleep cycles, and reproduction. The current study randomly chose 23 critical hypothalamic genes and measured the transcriptional levels (Table 5).

**Table 5.**
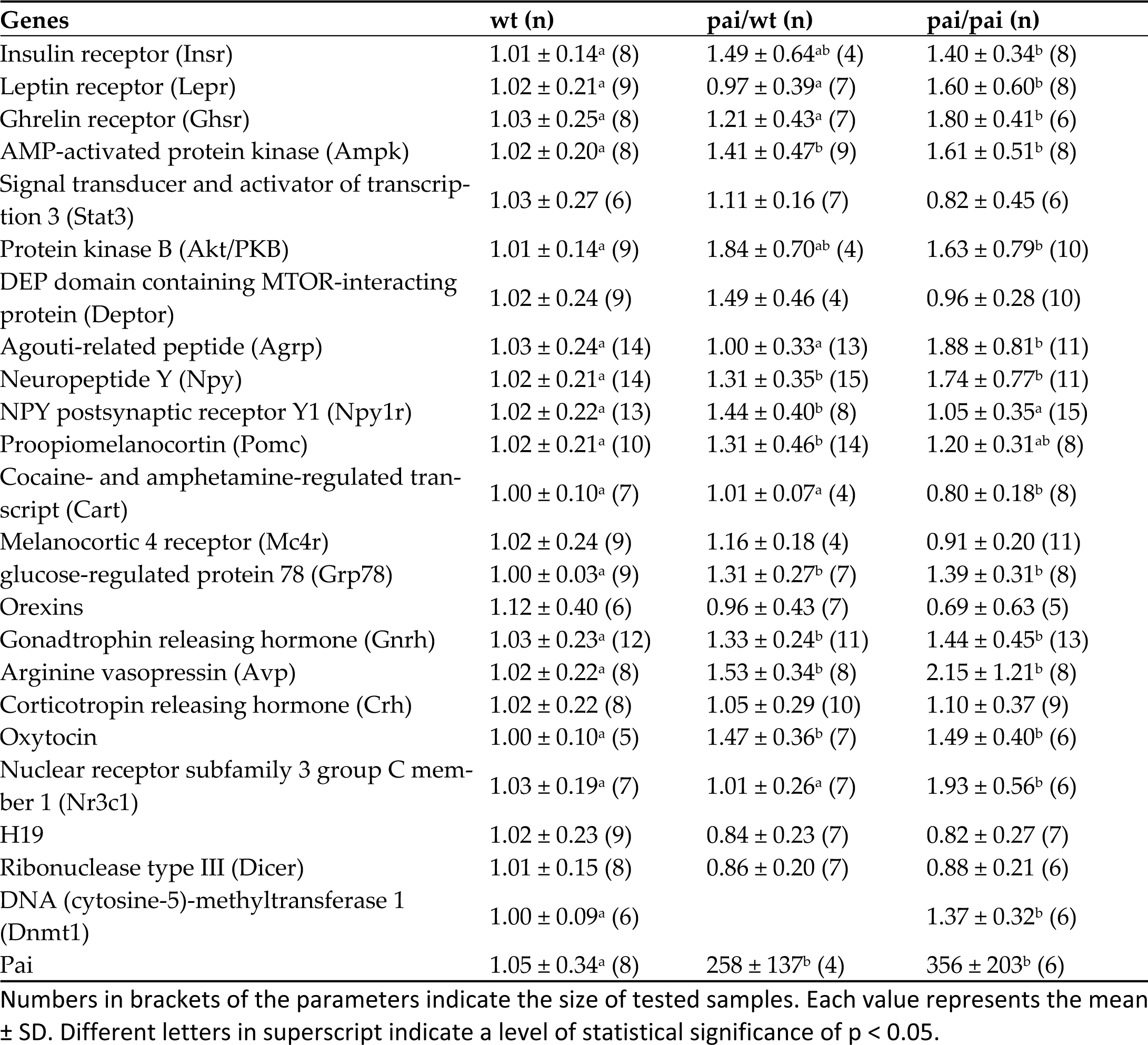
Relative expression of mRNAs in the hypothalamus.

Compared to wt samples, in the pai/pai hypothalamus, RNA level changes were observed in 14 (61%) genes. Only Cart, a marker of appetite-suppressive POMC neuron cells, was down-regulated among them. The remaining 13 genes were up-regulated, including appetite-relative genes such as the Insulin receptor, ghrelin receptor, agouti-related peptide, neuropeptide Y, leptin receptor, and the intracellularly signal molecule Akt/PKB, suggesting the sensitivity in regulating energy balance. In addition, the up-regulated gonadotrophin-releasing hormone, oxytocin, arginine vasopressin, DNA methyltransferase-1, and Nr3c1, suggesting that t10c12-CLA might affect the various neural circuitry in the hypothalamus. Furthermore, the RNA levels of Ampk and molecular chaperone Grp78 which releases the endoplasmic reticulum pressure and transmits thermogenesis signals to the BAT tissues were increased in the pai/pai hypothalamus. However, western blot analysis revealed that the protein levels of AMPK and pAMPK on Thr172 or Thr183 were not altered in the pai/pai hypothalamus compared with the wt samples (Fig. 8 A-B).

**Figure 8.**
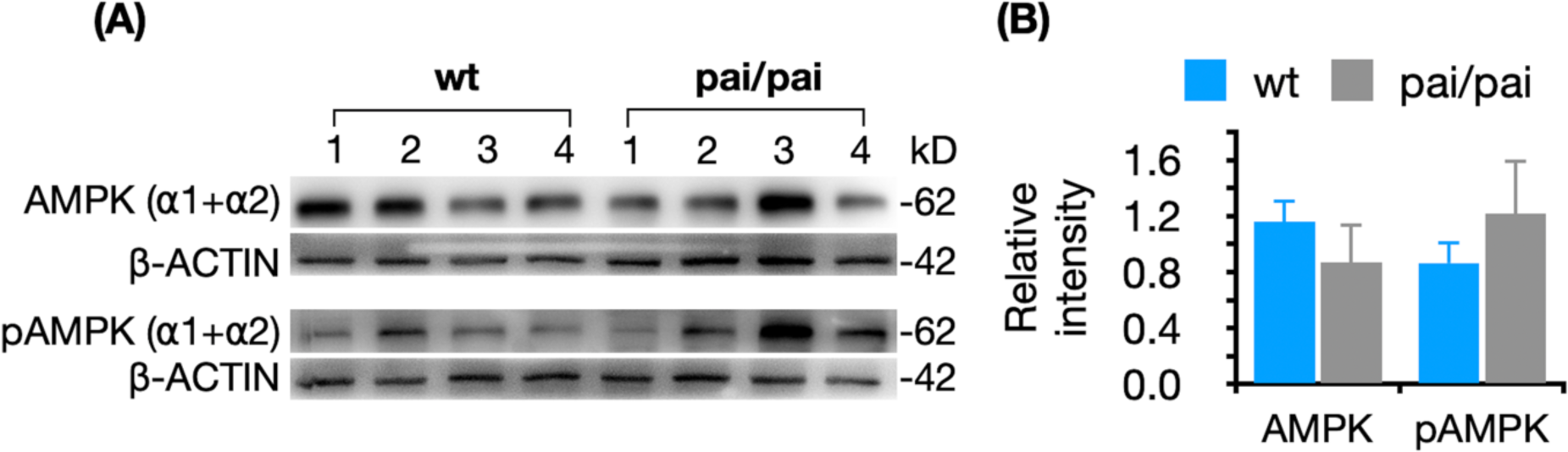
Western blot images (A) and relative intensities (B) analyses of the levels of AMPK (⍺1+⍺2) and pAMPK (⍺1-Thr183 + ⍺2-Thr172) in the hypothalamus from wt and pai/pai mice. Bars represent the mean ± SD.

In the pai/wt hypothalamus, the RNA levels of eight (36%) genes, including Ampk, Npy, Npy postsynaptic receptor Y1, Pomc, Grp78, GnRH, oxytocin, and arginine vasopressin, were increased, and no tested genes were down-regulated in pai/wt hypothalamus. Between pai/wt and pai/pai samples, the RNA levels of six (27%) genes, including Lepr, Ghsr, Agrp, Npy postsynaptic receptor Y1, Cart, and Nr3c1 exhibited differences (p < 0.05), suggesting the genotype-specific transcription in the hypothalamus.

## 4. Discussion

Unlike the temporary fat loss in the body [24] or milk fat depression which the fat de novo synthesis would reverse after the termination of the t10c12-CLA supplementation in mice [8], in pai mice, the chronic t10c12-CLA resulted in an irreversible fat reduction. It may cause an adaptive metabolic response throughout the entire developmental stage. Metabolic compensation or adaptation to t10c12-CLA is intrinsically complicated owing to its diverse metabolic functions in multiple target organs. Thus, interpreting these data could be more explicit in pai mice.

Although the fat reduction in milk did not impact the body weight of piglets from the energy supply perspective [25], previous studies in mice indicated that the dietary t10c12-CLA could reduce milk fat and retard pup growth [8,26]. However, we found that the wt pups fostered by the pai mother were reversely overweight at three weeks, suggesting that the natural t10c12-CLA from the mothers might play an active role during offspring development and more work needs to be done in future.

When a mouse was fed a low-fat diet containing t10c12-producing lactic acid bacteria or 0.4∼1% t10c12-CLA, its fat loss happened quickly, as early as seven days after starting the t10c12-CLA diet [13, 24, 27] and the body fat could reduce from approximately 2.8 g to less than 0.4 g in C57BL/6J males [28]. Simultaneously, the contents of t10c12-CLA were 0.1% in the livers of Balb/c males [7], approximately 0.2% in various tissues of ICR females [9], 0.45% in the milk of C57BL/6J females [8], 0.013 µmol/g in the brain or 0.089 µmol/g t10c12-CLA in eye lipids of C57BL/6N females [12]. Similarly, in the current study, the comparable values of t10c12-CLA were also detected in transgenic hearts (8.85 µg/g, 0.09%), livers (16.65 µg/g, 0.19%), kidneys (11.33 µg/g, 0.09%), BAT (28.8 µg/g, 0.23%), or WAT (4.89 µg/g, 0.13%; Supplementary Table 4) in pai mice.

However, it was difficult to explain why there were only a few minimal differences in the linoleic acid or its conjugate in various pai tissues. In our previous study in pai-transfected 3T3 cells, the drastic changes in an increase of product t10c12-CLA and a decrease of substrate linoleic acid could be observed easily [18]. In contrast, the present studies in the tissues had not shown the ideal changes. Frankly, we don’t know how to explain this discrepancy. Here we can only boldly speculate that linoleic acid is a relatively high component in somatic cells (10∼20 % in total fatty acids in organ/tissue homogenate) and the amount converted by PAI enzyme is limited.

Effect of dietary t10c12-CLA on food intake or appetite showed contradictory results in mice [4, 6, 27, 29]. In pai mice, we had not observed any changes in food intake within metabolic cages or crucial circulating leptin, insulin, glucose, and ghrelin. The results suggest that the anti-obesity property of t10c12-CLA is independent of food intake or appetite. However, the hypothalamic RNA increases in receptors for insulin, leptin, and ghrelin, and vital appetite-relative genes Npy, Argp, and an RNA decrease in appetite-suppressive Cart imply a tendency to promote food intake, which might be a mechanism of feedback regulation caused by more heat release, for the hormonal receptors, neuropeptides and energy sensor AMPK are in a complex regulation and interplay of energy homeostasis in the hypothalamic cells.

Investigations of oral t10c12-CLA administration suggested that the fat loss mainly occurred in fat depots through modulating adipocyte metabolism, such as increased fatty acid oxidation and browning WAT [2, 13, 30], or increasing the number of beige adipocytes in mice [28]. The pai/pai mice here had reduced WAT mass by approximately 20% at 11 weeks. However, RNA analysis indicated reduced pai/pai WAT browning and gluconeogenesis. Interestingly, the beta-oxidation of very long-chain FAs in peroxisomes was increased. That was, the pai/pai mice had adapted to long-term t10c12-induced fat reduction and could decrease the browning of WAT and mobilise the energy from the very long-chain FAs to maintain energy homeostasis.

Studies on whether the dietary t10c12-CLA modulates hepatic lipid metabolism and even induces fatty liver also showed contradictory results in mice [5, 13, 24, 31, 32]. It was supposed that the t10c12-induced hepatic steatosis might be due to the uptake and accumulation of lipids mobilised from the adipose tissue [5]. In the current study, although the essential genes Fasn, Dgat1, and Lpl involved in TGs uptake and synthesis were up-regulated in hepatocytes, analysis of hepatic TGs and TC levels and slices indicated no steatosis in pai livers. Furthermore, the blood TGs concentration was also lowered in pai mice, similar to the reduced plasma TGs in mice treated with a high dose of t10c12-CLA (0.6% w/w) [33]. It suggested that dietary t10c12-CLA might not induce fat accumulation in the liver when plenty of energy was consumed for lactation [8]. In pai mice, more energy expenditure via increased thermogenesis also hinted that no surplus energy could be accumulated in the liver. Our results suggest that the t10c12-CLA has yet to thoroughly break local lipid homeostasis in the livers.

A study in female 129Sv/J retired breeders indicated that dietary CLA mixture could induce body fat reduction via increasing energy expenditure [4]. Unfortunately, this work has yet to clarify whether the energy expenditure is via exercise strengthening or fat burning. Here, increased heat release and reduced activities in pai/pai mice suggest body fat reduction results from increased thermogenesis. However, the normal levels of circulating PGE2, adrenaline, corticosterone, TNF-⍺, Il-6, and CRP in pai mice suggest that the increased heat production is not due to the response to bacterial PAI enzyme-induced stress and/or inflammation in pai mice. That was, more heat release in pai mice resulted from the t10c12-CLA.

BAT thermogenesis can account for up to 50% of the body’s energetic maintenance demand. An increase [9] or decrease in BAT mass [33] were respectively observed in t10c12-CLA treated mice, but in pai/pai mice, the increased BAT mass might correlate with more heat release; additionally, the over-expression of UCP1 and UCP2 proteins in the pai/pai brown adipocytes also suggest increased thermogenesis via BAT activation. However, no change in the pAMPK levels suggests that the activated BAT heating and lipolysis are independent of the AMPK activation.

Based on the phenotypic characteristics of pai/wt mice, the authors suspected that the excess FGF21 in the blood and the central regulatory mechanism might be involved in the t10c12-CLA-induced BAT thermogenesis. FGF21 functions as a master sensitiser of specific hormonal signalling involved in enhancing insulin sensitivity and lowering serum glucose through direct actions on adipose tissues, possibly through the FGF21/adiponectin/ceramide axis [34], increasing energy expenditure through immediate efforts on the central nervous system to stimulate BAT thermogenesis, decreasing hepatic oxidative stress [35], lowering plasma TGs by accelerating lipoprotein catabolism in WAT and BAT [36], and regulating macronutrient preference [23]. In the current study, excess FGF21 in the blood suggests that the FGF21 plays an active role in BAT thermogenesis. However, the excess FGF21 appeared only in pai/wt, not in pai/pai mice suggests the t10c12-CLA’s impact on the body may be dose-dependent. Thus, the observable characteristics in pai/wt mice, such as an increase in FAs content in the kidney, body weight increase during adulthood, blood glucose sensitivity, serum TGs decrease, and genotype-specific gene expression in various tissues would be associated with the excess FGF21. We have yet to pay more attention to unique features in pai/wt mice here, and further work needs to be done in the future.

AMPK in the ventromedial nucleus of the hypothalamus can decrease ceramide-induced endoplasmic reticulum (ER) stress which stimulates the thermogenic program in BAT through the sympathetic nervous system [37]. In addition, GRP78 over-expression in the ventromedial nucleus can ameliorate hypothalamic ER stress, leading to weight loss, reduced hepatic steatosis, improved insulin and leptin sensitivities, increased BAT thermogenesis and stimulation WAT browning [38,39]. In the pai hypothalamus, the up-regulated transcriptional levels of Ampk and Grp78 suggest that the hypothalamic neurons might promote heat production via ameliorating ER stress in pai/pai mice. Unfortunately, we had not observed evident changes in the hypothalamic AMPK and pAMPK proteins.

Focused on increased thermogenesis, more work needs to clarify if the t10c12-CLA might affect ceramide in various ways: 1) the sphingomyelinase pathway to break down sphingomyelin in the plasma membrane to release ceramide; 2) de novo synthesis of ceramide from the condensation of palmitic acid (16:0) and serine to form 3-keto-dihydrosphingosine in the ER; or 3) the salvage pathway to break down sphingolipids to sphingosine in the lysosomes and to form ceramide by reacylation in the ventromedial nucleus.

According to this study’s metabolic profile in pai mice, we supposed that t10c12-CLA’s impact is genotype-specific (dose-dependent). That is, t10c12-CLA in low dose in pai/wt mice causes hepatic FGF21 secretion and excess FGF21 in the blood simultaneously increases the BAT thermogenesis and WAT reduction. Besides, t10c12-CLA in high doses in pai/pai mice directly acts on BAT to stimulate thermogenesis by promoting beta-oxidation and UCP1/2 thermogenesis, possible through immediate efforts on the central nervous system to induce the WAT loss (Fig. 9).

**Figure 9.**
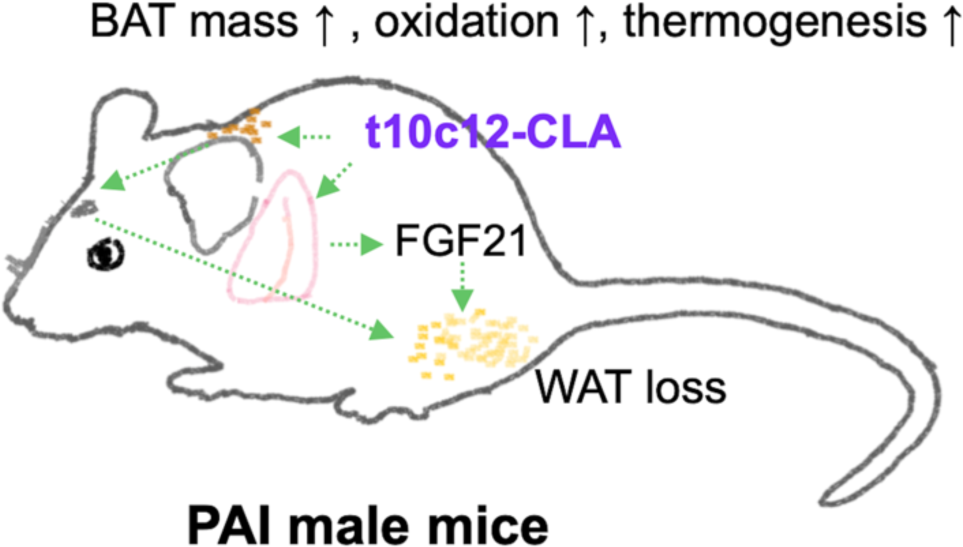
Schematic diagram illustrating t10c12-CLA action in a mouse. T10c12-CLA in low dose acts on the liver to stimulate the secretion of FGF21 which can promote BAT thermogenesis and WAT reduction. In addition, t10c12-CLA in high dose can also directly acts on BAT to induce its mass increase and promote thermogenesis, which can support immediate efforts on WAT reduction via central action. The black arrows indicate metabolic trends and the green arrows indicate possible action direction.

In addition, the expression modification of other hypothalamic genes, such as GCs receptor Nr3c1, required negative regulation of the hypothalamic–pituitary–adrenal axis at a basal condition and under stress [40]; vasopressin involved in fluid and electrolyte balance; GnRH and oxytocin involved in reproduction; even imprinted H19 and Dnmt1 involved in the epigenetic modification, were also observed in the pai hypothalamus. These aberrant expressions of hypothalamic genes suggest that the t10c12-CLA might play active roles in the myriad neural circuits in the hypothalamus and a more complex mechanism of action of t10c12-CLA needs to be investigated further.

## Abbreviations

CLA: conjugated linoleic acid
LA: linoleic acid
t10c12: *trans* 10, *cis* 12
pai: *Propionibacterium acnes* isomerase
wt: wild-type
FGF21: fibroblast growth factor 21
UCP: uncoupling protein
AMPK: AMP-activated protein kinase
BAT: brown adipose tissue
WAT: white adipose tissue
FAs: fatty acids

## Ethics approval and consent to participate

All animal experiments were performed following the Committee for Experimental Animals of Yangzhou University.

## Institutional Review Board Statement

The animal study protocol was approved by the Ethics Committee of Yangzhou University (protocol code NSFC2020-SYXY-20 and dated 25 March 2020).

## Consent for publication

Not applicable

## Availability of data and materials

The datasets used and/or analysed during the current study are available from the corresponding author upon reasonable request.

## Authors’ contributions

Conceptualization, Yu Rao, Shi Li, Sheng Cui and Ke Gou; Data curation, Yu Rao, Mei Li, Bao Wang, Yang Wang, Lu Liang, Shuai Yu and Zong Liu; Formal analysis, Yu Rao, Shi Li and Ke Gou; Funding acquisition, Yu Rao, Sheng Cui and Ke Gou; Investigation, Yu Rao, Mei Li, Bao Wang, Yang Wang and Lu Liang; Methodology, Yu Rao, Mei Li, Bao Wang, Yang Wang and Lu Liang; Project administration, Sheng Cui and Ke Gou; Software, Yu Rao, Shuai Yu and Zong Liu; Supervision, Ke Gou; Validation, Shuai Yu and Zong Liu; Visualization, Shuai Yu and Zong Liu; Writing – original draft, Yu Rao; Writing – review & editing, Ke Gou. All authors proofread, made comments, and approved the report.

## Acknowledgments

We thank Dr Li-Li Song and graduate students Yun-Cheng Zhuang, Yu-Huang Chen, Chao Wang, Qi Wang, Mei Liu, and Jia-Qi Lu in our lab for their generous assistance in partial tests and Ms Yu-Yang Wang in our university for technique help in gas chromatography. We also thank Professor Hong-Jun Wang at the Medical University of South Carolina, USA, for the helpful discussion during the writing process and Ms Wen-Jing Gou’s language help in preparing this manuscript.

## Supplementary data

This manuscript contains supplementary data.

## References

[1] Ritzenthaler KL, McGuire MK, McGuire MA, Shultz TD, Koepp AE, Luedecke LO, et al. Consumption of conjugated linoleic acid (CLA) from CLA-enriched cheese does not alter milk fat or immunity in lactating women. J Nutr. 2005;135:422–30.

[2] Park Y, Albright KJ, Liu W, Storkson JM, Cook ME, Pariza MW. Effect of conjugated linoleic acid on body composition in mice. Lipids. 1997;32:853–8.

[3] Baumgard LH, Corl BA, Dwyer DA, Saebo A, Bauman DE. Identification of the conjugated linoleic acid isomer that inhibits milk fat synthesis. Am J Physiol Regul Integr Comp Physiol. 2000;278:R179–84.

[4] Park Y, Park Y. Conjugated nonadecadienoic acid is more potent than conjugated linoleic acid on body fat reduction. J Nutr Biochem. 2010;21:764–73.

[5] Vyas D, Kadegowda AK, Erdman RA. Dietary conjugated linoleic acid and hepatic steatosis: species-specific effects on liver and adipose lipid metabolism and gene expression. J Nutr Metab. 2012;2012:932928.

[6] Kim JH, Kim Y, Kim YJ, Park Y. Conjugated linoleic acid: Potential health benefits as a functional food ingredient. Annu Rev Food Sci Technol. 2016;7:221–44.

[7] Rosberg-Cody E, Stanton C, O’Mahony L, Wall R, Shanahan F, Quigley EM, et al. Recombinant lactobacilli expressing linoleic acid isomerase can modulate the fatty acid composition of host adipose tissue in mice. Microbiol-Sgm. 2011;157:609–15.

[8] Harvatine KJ, Robblee MM, Thorn SR, Boisclair YR, Bauman DE. Trans-10, cis-12 CLA dose-dependently inhibits milk fat synthesis without disruption of lactation in C57BL/6J mice. J Nutr. 2014;144:1928–34.

[9] Li SL, Ma SY, Xu BR, Fan ZY, Li MJ, Cao WG, et al. Effects of trans-10, cis-12-conjugated linoleic acid on mice are influenced by the dietary fat content and the degree of murine obesity. Eur J Lipid Sci Tech. 2015;117:1908–18.

[10] Adkins Y, Belda BJ, Pedersen TL, Fedor DM, Mackey BE, Newman JW, et al. Dietary docosahexaenoic acid and trans-10, cis-12-conjugated linoleic acid differentially alter oxylipin profiles in mouse periuterine adipose tissue. Lipids. 2017;52:399–413.

[11] So MH, Tse IM, Li ET. Dietary fat concentration influences the effects of trans-10, cis-12 conjugated linoleic acid on temporal patterns of energy intake and hypothalamic expression of appetite-controlling genes in mice. J Nutr. 2009;139:145–51.

[12] Vemuri M, Adkins Y, Mackey BE, Kelley DS. Docosahexaenoic acid and eicosapentaenoic acid did not alter trans-10,cis-12 conjugated linoleic acid incorporation into mice brain and eye lipids. Lipids. 2017;52:763–9.

[13] Shen W, Baldwin J, Collins B, Hixson L, Lee KT, Herberg T, et al. Low level of trans-10, cis-12 conjugated linoleic acid decreases adiposity and increases browning independent of inflammatory signaling in overweight Sv129 mice. J Nutr Biochem. 2015;26:616–25.

[14] Marques TM, Wall R, O’Sullivan O, Fitzgerald GF, Shanahan F, Quigley EM, et al. Dietary trans-10, cis-12-conjugated linoleic acid alters fatty acid metabolism and microbiota composition in mice. Br J Nutr. 2015;113:728–38.

[15] Oteng AB, Kersten S. Mechanisms of action of trans fatty acids. Adv Nutr. 2020;11:697–708.

[16] Kohno-Murase J, Iwabuchi M, Endo-Kasahara S, Sugita K, Ebinuma H, Imamura J. Production of trans-10, cis-12 conjugated linoleic acid in rice. Transgenic Res. 2006;15:95–100.

[17] Wang C, Wang YY, Rao Y, Li MJ, Liu ZP, Li SL, et al. Heterologous expression of Propionibacterium acnes isomerase in mouse (Mus musculus) cells and production of t10c12-conjugated linoleic acid. Chinese Journal of Agricultural Biotechnology. 2021;29:2304–11.

[18] Li SL. Analysis of the effect of trans 10, cis 12-conjugated linoleic acid on mice growth and metabolism. Ph.D. Dissertation. Beijing: China Agricultural University; 2015.

[19] Li M, Li S, Rao Y, Cui S, Gou K. Loss of smooth muscle myosin heavy chain results in the bladder and stomach developing lesion during foetal development in mice. J Genet. 2018;97:469–75.

[20] Chu VT, Weber T, Graf R, Sommermann T, Petsch K, Sack U, et al. Efficient generation of Rosa26 knock-in mice using CRISPR/Cas9 in C57BL/6 zygotes. BMC Biotechnol. 2016;16:4.

[21] Jenkins TC. Technical note: common analytical errors yielding inaccurate results during analysis of fatty acids in feed and digesta samples. J Dairy Sci. 2010;93:1170–4.

[22] Chen HC, Farese RV, Jr. Determination of adipocyte size by computer image analysis. J Lipid Res. 2002;43:986–9.

[23] Flippo KH, Potthoff MJ. Metabolic Messengers: FGF21. Nat Metab. 2021;3:309–17.

[24] Cordoba-Chacon J, Sugasini D, Yalagala PCR, Tummala A, White ZC, Nagao T, et al. Tissue-dependent effects of cis-9,trans-11-and trans-10,cis-12-CLA isomers on glucose and lipid metabolism in adult male mice. J Nutr Biochem. 2019;67:90–100.

[25] Sandri EC, Harvatine KJ, Oliveira DE. Trans-10, cis-12 conjugated linoleic acid reduces milk fat content and lipogenic gene expression in the mammary gland of sows without altering litter performance. Br J Nutr. 2020;123:610–8.

[26] Robblee MM, Boisclair YR, Bauman DE, Harvatine KJ. Dietary fat does not overcome trans-10, cis-12 conjugated linoleic acid inhibition of milk fat synthesis in lactating mice. Lipids. 2020;55:201–12.

[27] Shelton VJ, Shelton AG, Azain MJ, Hargrave-Barnes KM. Incorporation of conjugated linoleic acid into brain lipids is not necessary for conjugated linoleic acid-induced reductions in feed intake or body fat in mice. Nutr Res. 2012;32:827–36.

[28] Yeganeh A, Zahradka P, Taylor CG. Trans-10,cis-12 conjugated linoleic acid (t10-c12 CLA) treatment and caloric restriction differentially affect adipocyte cell turnover in obese and lean mice. Journal of Nutritional Biochemistry. 2017;49:123–32.

[29] Park Y, Albright KJ, Storkson JM, Liu W, Pariza MW. Conjugated linoleic acid (CLA) prevents body fat accumulation and weight gain in an animal model. J Food Sci. 2007;72:S612–7.

[30] den Hartigh LJ, Wang SR, Goodspeed L, Wietecha T, Houston B, Omer M, et al. Metabolically distinct weight loss by 10,12 CLA and caloric restriction highlight the importance of subcutaneous white adipose tissue for glucose homeostasis in mice. Plos One. 2017;12.

[31] Ashwell MS, Ceddia RP, House RL, Cassady JP, Eisen EJ, Eling TE, et al. Trans-10, cis-12-conjugated linoleic acid alters hepatic gene expression in a polygenic obese line of mice displaying hepatic lipidosis. J Nutr Biochem. 2010;21:848–55.

[32] Kostogrys RB, Franczyk-Zarow M, Maslak E, Gajda M, Mateuszuk L, Chlopicki S. Effects of margarine supplemented with t10c12 and C9T11 CLA on atherosclerosis and steatosis in apoE/LDLR -/- mice. J Nutr Health Aging. 2012;16:482–90.

[33] Shen W, Chuang CC, Martinez K, Reid T, Brown JM, Xi L, et al. Conjugated linoleic acid reduces adiposity and increases markers of browning and inflammation in white adipose tissue of mice. J Lipid Res. 2013;54:909–22.

[34] Holland WL, Adams AC, Brozinick JT, Bui HH, Miyauchi Y, Kusminski CM, et al. An FGF21-adiponectin-ceramide axis controls energy expenditure and insulin action in mice. Cell Metab. 2013;17:790–7.

[35] Ye D, Wang Y, Li H, Jia W, Man K, Lo CM, et al. Fibroblast growth factor 21 protects against acetaminophen-induced hepatotoxicity by potentiating peroxisome proliferator-activated receptor coactivator protein-1alpha-mediated antioxidant capacity in mice. Hepatology. 2014;60:977–89.

[36] Schlein C, Talukdar S, Heine M, Fischer AW, Krott LM, Nilsson SK, et al. Fgf21 lowers plasma triglycerides by accelerating lipoprotein catabolism in white and brown adipose tissues. Cell Metab. 2016;23:441–53.

[37] Martinez-Sanchez N, Seoane-Collazo P, Contreras C, Varela L, Villarroya J, Rial-Pensado E, et al. Hypothalamic ampk-er stress-jnk1 axis mediates the central actions of thyroid hormones on energy balance. Cell Metab. 2017;26:212–29 e12.

[38] Contreras C, Gonzalez-Garcia I, Seoane-Collazo P, Martinez-Sanchez N, Linares-Pose L, Rial-Pensado E, et al. Reduction of hypothalamic endoplasmic reticulum stress activates browning of white fat and ameliorates obesity. Diabetes. 2017;66:87–99.

[39] Contreras C, Fondevila MF, Lopez M. Hypothalamic GRP78, a new target against obesity? Adipocyte. 2018;7:63–6.

[40] Laryea G, Schutz G, Muglia LJ. Disrupting hypothalamic glucocorticoid receptors causes HPA axis hyperactivity and excess adiposity. Mol Endocrinol. 2013;27:1655–65.

