## Supplemental files for "Metabolic impact of trans 10, cis 12-conjugated linoleic acid on pai transgenic mice"

**Supplementary Table 1. Sources of dietary ingredients.**

| Ingredients | Grams | Sources |
|---|---|---|
| Lactic casein | 200.0 | Gansu Hualing Dairy Co. Ltd, Hezuo city, China |
| L-cystine | 3.0 | Zhongtuan shihua Food Reagents Co. Ltd, Hangzhou, China |
| Sucrose | 176.8 | Jinhuitaiya, Chemcal Reagents Co. Ltd, Tianjin, China |
| Corn Starch | 452.2 | Zhucheng Xingmao Corn Co. Ltd, Weifang city, China |
| Maltodextrin 10 | 75.0 | Dongzhu Food Co. Ltd, Guilin city, China |
| Cellulose | 50.0 | Yisheng Kangyuan Biotech Co. Ltd, Zibo city, China |
| Soybean oil | 25.0 | Shandong Luhua Co. Ltd, Leiyang, China |
| Lard | 20.0 | Yaodong Baofengyuan Co. Ltd, Qingdao, China |
| Mineral Mix (20x) | 50.0 | Mingxin Chemical Reagents Co. Ltd, Zhenzhou, China |
| Choline bitartrate | 2.0 | Zhejiang Linuo Biotech Co. Ltd, Yiwu, China |
| Vitamin Mix (1000x) | 1.0 | Mingxin Chemical Reagents Co. Ltd, Zhenzhou, China |
| Total Gram | 1055 | |

**Supplementary Table 2. Primers sequences in transgene analysis**

| Gene | | Fragment length (bp) | Forward (top) and reverse (bottom) primers (5' to 3') | Annealing temperature (°C) |
|---|---|---|---|---|
| **PCR:** | | | | |
| | pai | 514 | TAACCATGTTCATGCCTTCTTC;<br>CACCTTGTTGTAGTGTCCGTTT | 57 |
| | Rosa26 | 605 | CCAAAGTCGCTCTGAGTTGTTATCAGT;<br>GGAGCGGGAGAAATGGATATGAAG | 57 |
| **Southern blot:** | | | | |
| | pai | 678 | CAGGATTCCACGATTACAC；<br>CAGTCAGGACCAGCACAT | 61 |
| **RT-PCR:** | | | | |
| | pai | 164 | GGATTCCACGATTACACCA;<br>GGGTCATCAGCCCATCTA | 60 |
| | Gaphd | 78 | CCTGCTGGATATCATTAAAGCACTG;<br>GTCAAGGGCATATCCAACAACAAAC | 60 |

**Supplementary Table 3. Primer sequences of real-time PCR.**

| mRNA | Full name | Forward (top) and reverse (bottom) primers (5’ to 3’) | Genbank accession no. |
|---|---|---|---|
| *36B4* | Ribosomal protein, large, P0 | CACTGGTCTAGGACCCGAGAAG; GGTGCCTCTGGAGATTTTCG | NM_007475.5 |
| *Acaa2* | Acetyl-CoA acyltransferase 2 | CTGCTACGAGGTGTGTTCATC; AGCTCTGCATGACATTGCCC | NM_177470.3 |
| *Acbp* | Acyl-CoA-binding protein | CATCCGTATCACCTCACC; GTATTTACATCGCCCACA | NM_007830.4 |
| *Acox1* | Acyl-CoA oxidase 1, transcript variant 1 | AGATTGGTAGAAATTGCTGCAAAA; ACGCCACTTCCTTGCTCTTC | NM_015729.3 |
| *Acox2* | Acyl-CoA oxidase 2, transcript variant 1 | AACCCAGGGGATCGAGTGT; CGCAGCTCAGTGTTTGGGAT | NM_053115.2 |
| *Adcy3* | Adenylate cyclase 3, transcript variant 1 | AGATGTTCGGTGCCACCTG; ACTTCACCAGGGCTTCGTAAG | NM_138305.3 |
| | Adiponectin | AGCCGCTTATATGTATCGCTCA； TGCCGTCATAATGATTCTGTTGG | NM_009605.5 |
| *Agpat1* | 1-acylglycerol-3-phosphate O-acyltransferase 1 | GCTGGCTGGCAGGAATCAT; GTCTGAGCCACCTCGGACAT | NM_018862 |
| *Agpat2* | 1-acylglycerol-3-phosphate O-acyltransferase 2 | TTTGAGGTCAGCGGACAGAA; AGGATGCTCTGGTGATTAGAGATGA | NM_026212 |
| *Agrp* | Agouti related neuropeptide | GGCCTCAAGAAGACAACTGC; GACTCGTGCAGCCTTACACA | NM_007427.3 |

| mRNA | Full name | Forward (top) and reverse (bottom) primers (5' to 3') | Genbank accession no. |
|---|---|---|---|
| *Ampk* | AMP-activated protein kinase | TTGACGATGAGGCTGTGAAG;<br>ATAAGCCACTGCAAGCTGGT | NM_178143.2 |
| *ApoB* | Apolipoprotein B | CGTGGGCTCCAGCATTCTA;<br>TCACCAGTCATTTCTGCCTTTG | NM_009693.2 |
| *Atgl* | Adipose triglyceride lipase | TTCACCATCCGCTTGTTGGAG;<br>AGATGGTCACCCAATTTCCTC | NM_025802.3 |
| *Avp* | Arginine vasopressin | GCTCAACACTACGCTCTC；<br>CTTGGGCAGTTCTGGAAG | NM_009732.2 |
| *Cart* | Cocaine- and amphetamine-regulated transcript | GTCCCACGAGAAGGAGCTGCCAA;<br>GCCCATCCGCTCTCTGAGGGG | NM_013732.7 |
| *Cd11c* | CD11 antigen-like family member C, integrin alpha X | CTGGATAGCCTTTCTTCTGCTG;<br>GCACACTGTGTCCGAACTC | NM_021334.3 |
| *Cd36* | CD36 antigen, transcript variant 1 | GGACATTGAGATTCTTTTCCTCTG;<br>GCAAAGGCATTGGCTGGAAGAAC | NM_00115955<br>8.1 |
| *Cd68* | CD68 antigen | CTTCCCACAGGCAGCACAG;<br>AATGATGAGAGGCAGCAAGAGG | NM_00129105<br>8.1 |
| *Cebpβ* | CCAAT/enhancer binding protein (C/EBP), beta | CTGCGGGGTTGTTGATGT;<br>ATGCTCGAAACGGAAAAGGT | NM_00128773<br>8.1 |
| *Cgi-58* | Comparative gene identification 58 | TGGATTCTTGGCTGCTGCTTAC;<br>TTAAAGGGAGTCAATGCTGCTC | NM_026179.2 |
| *Chrebp* | Carbohydrate response element binding protein | CCTTCGCCAACTCAGCACTT;<br>TGGCTTGCTCAGGCACAA | NM_021455 |
| *Cpt1a* | Carnitine palmitoyltransferase 1a | CACCAACGGGCTCATCTTCTA;<br>CAAAATGACCTAGCCTTCTATCGAA | NM_013495 |
| *Crh* | Corticotropin releasing hormone | GGCATCCTGAGAGAAGTCCC；<br>GTTAGGGGCGCTCTCTTCTC | NM_205769.3 |

| mRNA | Full name | Forward (top) and reverse (bottom) primers (5' to 3') | Genbank accession no. |
|---|---|---|---|
| *Cypb* | Cyclophilin B | TGGAGAGCACCAAGACAGACA; TGCCGGAGTCGACAATGAT | NM_011149.2 |
| *Deptor* | DEP domain containing MTOR-interacting protein | ATAGACGGCACCATCTCAAAAC; GTCGGCTAATTTCTGCATGAGT | NM_145470.3 |
| *Dgat1* | Diacylglycerol acyltransferase 1 | GAGGCCTCTCTGCCCCTATG; GCCCCTGGACAACACAGACT | NM_010046 |
| *Dgat2* | Diacylglycerol acyltransferase 2 | CCGCAAAGGCTTTGTGAAG; GGAATAAGTGGGAACCAGATCA | NM_026384 |
| *Dicer* | Ribonuclease type III | CACACGCCTCCTACCACTACAACAC; CCGTGGGTCTTCATAAAGGT | NM_148948.2 |
| *Dnmt1* | DNA (cytosine-5)-methyltransferase 1 | GTCGGACAGTGACACCCTTT; TGGGTTTCCGTTTAGTGGGG | NM_00119943 1.1 |
| *F4/80* | Adhesion G protein-coupled receptor E1 | CTTTGGCTAAGGGCTTCCAGTC; GCAAGGAGGACAGAGTTTATCGTG | NM_010130.4 |
| *Fads1* | Fatty acid desaturase 1 | AGCACATGCCATACAACCATC; TTTCCGCTGAACCACAAAATAGA | NM_146094.2 |
| *Fads2* | Fatty acid desaturase 2 | GATGGCTGCAACATGACTATGG; GCTGAGGCACCCTTTAAGTGG | NM_019699.2 |
| *Fasn* | Fatty acid synthase | GCTGCGGAAACTTCAGGAAAT; AGAGACGTGTCACTCCTGGACTT | NM_007988.3 |
| *Fgf21* | Fibroblast growth factor 21 | CTGCTGGGGGTCTACCAAG; CTGCGCCTACCACTGTTCC | NM_020013.4 |
| *Foxa2* | Forkhead box protein a2 | ACTTTGGGAGAGCTTTGAGGAA; CCCATCTATTTAGGGACACAGACA | NM_010446.3 |
| *Foxc2* | Forkhead box protein C2 | AAAGCGCCCCTCTCTCAGA; CTCAAACTGAGCTGCGGATAAGT | NM_013519.2 |

| mRNA | Full name | Forward (top) and reverse (bottom) primers (5' to 3') | Genbank accession no. |
|---|---|---|---|
| *G6p* | Glucose-6-phosphatase | TGGGCAAAATGGCAAGGA; TCTGCCCCAGGAATCAAAAAT | NM_008061.4 |
| *G6pd* | Glucose-6-phosphate dehydrogenase | GAACGCAAAGCTGAAGTGAGACT; TCATTACGCTTGCACTGTTGGT | NM_019468.2 |
| *Gapdh* | Glyceraldehyde-3-phosphate de-hydrogenase, transcript variant 1 | GAACATCATCCCTGCATCC; CCAGTGAGCTTCCCGTTCA | NM_001289726.1 |
| *Gck* | Glucokinase | CCGTGATCCGGGAAGAGAA; GGGAAACCTGACAGGGATGAG | NM_010292.5 |
| *Ghsr* | ghrelin receptor | GGACCAGAACCACAAACAGACA; CAGCAGAGGATGAAAGCAAACA | NM_177330.4 |
| *Glut4* | Glucose transporter type 4, transcript variant 1 | GTGACTGGAACACTGGTCCTA; CCAGCCACGTTGCATTGTAG | NM_009204.2 |
| *Gnrh* | Gonadotropin releasing hormone 1, transcript variant 1 | CACTGGTCCTATGGGTTGC; TTCTGCCTGGCTTCCTCT | NM_008145.3 |
| *Gpat1* | Glycerol-3-phosphate acyltrans-ferase 1, transcript variant 1 | CAACACCATCCCCGACATC; GTGACCTTCGATTATGCGATCA | NM_001356285.1 |
| *Grp78* | Glucose-regulated protein 78, transcript variant 1 | ACTTGGGGACCACCTATTCCT; ATCGCCAATCAGACGCTCC | NM_001163434.1 |
| *H19* | H19, transcript variant 1 | GGAATGTTGAAGGACTGAGGG; GTAACCGGGATGAATGTCTGG | NR_130973.1 |
| *Hmgcr* | HMG-CoA reductase, transcript variant 1 | CTTGTGGAATGCCTTGTGATTG; AGCCGAAGCAGCACATGAT | NM_008255.2 |
| *Hsl* | Hormone-sensitive lipase, transcript variant 1 | GGAGCACTACAAACGCAACGA; TCGGCCACCGGTAAAGAG | NM_010719.5 |
| *Htgl* | Hepatic triglyceride lipase, transcript variant 1 | ATGGGAAATCCCCTCCAAATCT; GTGCTGAGGTCTGAGACGA | NM_008280.2 |

| mRNA | Full name | Forward (top) and reverse (bottom) primers (5’ to 3’) | Genbank accession no. |
|---|---|---|---|
| *Igf1* | Insulin-like growth factor 1, transcript variant 1 | GCCCCACTGAAGCCTACAAA; TGAGTCTTGGGCATGTCAGTGT | NM_010512.5 |
| *Igf2r* | Insulin-like growth factor 2 receptor | GCCTCTGGAGACATGAGGAC; GGTGCCTACTTTCCTGCTGA | NM_010515.2 |
| *Igfbp1* | Insulin-like growth factor binding protein 1 | ATCAGCCCATCCTGTGGAAC; TGCAGCTAATCTCTCTAGCACTT | NM_008341.4 |
| *Insig1* | Insulin induced gene 1 | TCACAGTGACTGAGCTTCAGCA; TCATCTTCATCACACCCAGGAC | NM_153526.5 |
| *Insig2a* | Insulin induced gene 2a, transcript variant 2 | CCCTCAATGAATGTACTGAAGGATT; TGTGAAGTGAAGCAGACCAATGT | NM_178082.3 |
| *Insr* | Insulin receptor, transcript variant 1 | CGAGTGCCCGTCTGGCTATA; GGCAGGGTCCCAGACATG | NM_010568.3 |
| *Irs1* | Insulin receptor substrate-1 | TCACAGCAGAATGAAGACC; CTACTGATGAGGAAGATATGAGG | NM_010570.4 |
| *Irs2* | Insulin receptor substrate-2 | GGAGAACCCAGACCCTAAGCTACT; GATGCCTTTGAGGCCTTCAC | NM_001081212.2 |
| *Lcad* | Long-chain acyl-CoA dehydrogenase | TCAATGGAAGCAAGGTGTTCA; GCCACGACGATCACGAGAT | NM_007381.4 |
| *Lchad* | Long-chain 3-hydroxyacyl-CoA dehydrogenase | TGCATTTGCCGCAGCTTTAC; GTTGGCCCAGATTTCGTTCA | NM_178878.3 |
| *Ldlr* | Low density lipoprotein receptor, transcript variant 1 | AGGCTGTGGGCTCCATAGG; TGCGGTCCAGGGTCATCT | NM_010700.3 |
| *Lepr* | Leptin receptor, transcript variant 1 | TGGTCCCAGCAGCTATGGT; ACCCAGAGAAGTTAGCACTGT | NM_146146.3 |
| *Lkb1* | Liver kinase B1, transcript variant 2 | GACTTCACAGTGCCTGGTGTC; AGCAGACAGGGAGCTACACTA | NM_001301853.2 |

| mRNA | Full name | Forward (top) and reverse (bottom) primers (5' to 3') | Genbank accession no. |
|---|---|---|---|
| *Lpl* | Lipoprotein lipase | ACTATGTGTCTAACTGCCACTTCAA;<br>ATACATTCCCGTTACCGTCCAT | NM_008509.2 |
| *Lxrα* | Liver X receptor α, transcript variant 1 | TACGTCTCCATCAACCACCCC;<br>ACTTGCTCTGAATGGACGCTG | NM_013839.4 |
| *Lxrβ* | Liver X receptor β, transcript variant 1 | GCAGTTGGCACTAGAAG;<br>GGTAGGCTGAGGTGTAA | NM_009473.3 |
| *Malic* | Malic enzyme 1 | GCCGGCTCTATCCTCCTTTG;<br>TTTGTATGCATCTTGCACAATCTTT | NM_008615.2 |
| *Mc4r* | Melanocortic 4 receptor | CCCGGACGGAGGATGCTAT;<br>TCGCCACGATCACTAGAATGT | NM_016977.4 |
| *Mcad* | Medium-chain acyl-CoA dehydrogenase | GCAACTGCCCGCAAGTTT;<br>TACTCCCCGCTTTTGTCATATTC | NM_007382.5 |
| *Mgl* | Monoglyceride lipase, transcript variant 1 | CGGACTTCCAAGTTTTTGTCAGA;<br>GCAGCCACTAGGATGGAGATG | NM_00116625<br>1.1 |
| *Npy* | Neuropeptide Y | CCTTCCATGTGGTGATGGGA;<br>GCAGACTGGTTTCAGGGGAT | NM_023456.3 |
| *Npy1r* | Neuropeptide Y postsynaptic receptor Y1, transcript variant 1 | CACAGGCTGTCTTACACG;<br>GCGAATGTATATCTTGAAGTAG | NM_010934.4 |
| *Nr3c1* | Nuclear receptor subfamily 3 group C member 1, transcript variant 1 | GGAAGCGTGATGGACTTGTAT;<br>GCTTGGAATCTGCCTGAGAA | NM_008173.4 |
| *Orexins* | Orexins-total | GTCGCCAGAAGACGTGTTC;<br>GGTGGTAGTTACGGTCGGAC | NM_010410.2 |
| *Oxt* | Oxytocin | TGGCTTACTGGCTCTGACCT;<br>GGCAGGTAGTTCTCCTCCTG | NM_011025.4 |
| *Pai* | P*ropionibacterium acnes* isomerase | TGACGAGCGGGAATACTTTA；<br>GAGGGTCATCAGCCCATCTA | |

| mRNA | Full name | Forward (top) and reverse (bottom) primers (5’ to 3’) | Genbank accession no. |
|---|---|---|---|
| *Pepck* | Phosphoenolpyruvate carboxykinase | CCACAGCTGCTGCAGAACA; GAAGGGTCGCATGGCAAA | NM_011044.3 |
| *Pgc1α* | Ppar-γ coactivator 1 alpha, transcript variant 1 | CCCTGCCATTGTTAAGACC; TGCTGCTGTTCCTGTTTTC | NM_008904.3 |
| *Pgd* | 6-Phosphogluconate dehydrogenase, transcript variant 1 | TGAAGGGTCCTAAGGTGGTCC; CCGCCATAATTGAGGGTCCAG | NM_001081274.2 |
| *Pgr* | Progesterone receptor | CTCCGGGACCGAACAGAGT；ACAACAACCCTTTGGTAGCAG | NM_008829.2 |
| *Pi3k* | Phosphoinositide 3-kinase, transcript variant 2 | GACAGCGAAGCGACGGC; GTCTGATTTTACTGCCACGCTC | NM_001077495.2 |
| *Plin1* | Perilipin 1a | CTGTGTGCAATGCCTATGAGA; CTGGAGGGTATTGAAGAGCCG | NM_175640.2 |
| *Pomc* | Proopiomelanocortin, transcript variant 1 | CTCCTGCTTCAGACCTCCAT; CAGTCAGGGGCTGTTCATCT | NM_001278581.1 |
| *Ppar-γ* | Peroxisome proliferator-activated receptor-γ, transcript variant 1 | CACAATGCCATCAGGTTTGG; GCTGGTCGATATCACTGGAGATC | NM_001127330.2 |
| *Prdm16* | PR domain containing 16 | GACTTGGACACTACCACGGG; AGATGCACCCCCAAACTCAG | NM_027504.3 |
| *Scap* | SREBP cleavage-activating protein | ATTTGCTCACCGTGGAGATGTT; GAAGTCATCCAGGCCACTACTAATG | NM_001001144.3 |
| *Scd1* | Stearoyl-coA desaturase 1 | TCCTCCCTACCTCCAACT; CAACAACCAACCCTCGCA | NM_009127.4 |
| *Srebp1a* | Sterol regulatory element binding protein 1a, transcript variant X1/2 | GGCCGAGATGTGCGAACT; TTGTTGATGAGCTGGAGCATGT | NM_011480.4 |
| *Srebp1c* | Sterol regulatory element binding protein 1c, transcript variant X3 | GGAGCCATGGATTGCACATT; GGCCCGGGAAGTCACTGT | NM_001358314.1 |

| mRNA | Full name | Forward (top) and reverse (bottom) primers (5' to 3') | Genbank accession no. |
|---|---|---|---|
| *Srebp2* | Sterol regulatory element binding protein 2 | GCGTTCTGGAGACCATGGA; ACAAAGTTGCTCTGAAAACAAATCA | NM_033218.1 |
| *Stat3* | Signal transducer and activator of transcription 3 | CTGTAGAGCCATACACCAAGCAGCAGC; GGTCTTCAGGTACGGGGCAGCAC | NM_213659.3 |
| *Ucp1* | Uncoupling protein 1 | GAGGTGTGGCAGTGTTCATTG; GGCTTGCATTCTGACCTTCA | NM_009463.3 |
| *Ucp2* | Uncoupling protein 2 | GCTTCTGCACCACCGTCAT; GCCCAAGGCAGAGTTCATGT | NM_011671.5 |

Supplemental Table 4. Fatty acids (mg/g) in wt and pai tissues

| | wt | pai/wt | pai/pai | wt | pai/wt | pai/pai |
|---|---|---|---|---|---|---|
| ***Hearts*** | | | | ***Kidneys*** | | |
| No. of samples | 11 | 8 | 8 | 10 | 13 | 8 |
| 8:0 | 0.01 ± 0.00 | 0.01 ± 0.00 | 0.01 ± 0.00 | 0.01 ± 0.00$^{a}$ | 0.01 ± 0.00$^{b}$ | 0.01 ± 0.00$^{a}$ |
| 10:0 | ND | ND | ND | ND | ND | ND |
| 12:0 | ND | ND | ND | ND | ND | ND |
| 14:0 | 0.01 ± 0.00 | 0.01 ± 0.00 | 0.02 ± 0.00 | 0.02 ± 0.00$^{a}$ | 0.04 ± 0.02$^{b}$ | 0.02 ± 0.01$^{a}$ |
| 14:1 | ND | ND | ND | 0.01 ± 0.00$^{a}$ | 0.01 ± 0.00$^{b}$ | 0.01 ± 0.00$^{a}$ |
| 16:0 | 1.36 ± 0.07$^{a}$ | 1.44 ± 0.15$^{ab}$ | 1.60 ± 0.16$^{b}$ | 1.87 ± 0.21$^{a}$ | 2.56 ± 0.44$^{b}$ | 2.10 ± 0.23$^{c}$ |
| 16:1n-7 | 0.05 ± 0.02 | 0.06 ± 0.01 | 0.06 ± 0.01 | 0.06 ± 0.01$^{a}$ | 0.16 ± 0.07$^{b}$ | 0.08 ± 0.03$^{a}$ |
| 18:0 | 1.90 ± 0.24$^{ab}$ | 1.83 ± 0.23$^{a}$ | 2.07 ± 0.19$^{b}$ | 1.67 ± 0.19$^{a}$ | 2.06 ± 0.44$^{b}$ | 1.85 ± 0.17$^{b}$ |
| trans-18:1 | 0.02 ± 0.01 | 0.03 ± 0.02 | 0.03 ± 0.01 | 0.01 ± 0.00$^{a}$ | 0.01 ± 0.00$^{b}$ | 0.01 ± 0.00$^{a}$ |
| 18:1n-9 | 0.77 ± 0.10$^{a}$ | 0.79 ± 0.12$^{a}$ | 0.94 ± 0.12$^{b}$ | 0.79 ± 0.14$^{a}$ | 1.15 ± 0.27$^{b}$ | 0.90 ± 0.10$^{a}$ |
| 18:1n-7 | 0.29 ± 0.03$^{a}$ | 0.33 ± 0.05$^{ab}$ | 0.35 ± 0.03$^{b}$ | 0.29 ± 0.04$^{a}$ | 0.46 ± 0.10$^{b}$ | 0.33 ± 0.03$^{c}$ |
| 18:2n-6 | 2.06 ± 0.31 | 2.03 ± 0.31 | 2.11 ± 0.19 | 1.13 ± 0.16$^{a}$ | 1.38 ± 0.26$^{b}$ | 1.20 ± 0.19$^{a}$ |
| 20:0 | 0.01 ± 0.00 | 0.01 ± 0.00 | 0.01 ± 0.00 | 0.01 ± 0.00$^{a}$ | 0.02 ± 0.00$^{b}$ | 0.01 ± 0.00$^{b}$ |
| 18:3n-6 | ND | ND | ND | 0.01 ± 0.00$^{a}$ | 0.01 ± 0.00$^{b}$ | 0.01 ± 0.00$^{ab}$ |
| 18:3n-3 | 0.01 ± 0.00 | 0.01 ± 0.00 | 0.01 ± 0.00 | 0.01 ± 0.00$^{a}$ | 0.01 ± 0.00$^{b}$ | 0.01 ± 0.00$^{a}$ |
| t10c12-CLA | 0.007 ± 0.00 | 0.009 ± 0.00 | 0.008 ± 0.00 | 0.007 ± 0.00$^{a}$ | 0.011 ± 0.00$^{b}$ | 0.007 ± 0.00$^{a}$ |
| t10c12-CLA (%) | 0.07 ± 0.00$^{a}$ | 0.09 ± 0.01$^{b}$ | 0.08 ± 0.02$^{ab}$ | 0.07 ± 0.01$^{a}$ | 0.09 ± 0.02$^{b}$ | 0.06 ± 0.01$^{a}$ |
| 20:3n-6 | 0.13 ± 0.03 | 0.12 ± 0.02 | 0.12 ± 0.01 | 0.13 ± 0.04 | 0.15 ± 0.04 | 0.13 ± 0.04 |
| 20:4n-6 | 1.26 ± 0.24 | 1.23 ± 0.16 | 1.24 ± 0.12 | 2.05 ± 0.31$^{a}$ | 2.53 ± 0.65$^{b}$ | 2.21 ± 0.25$^{ab}$ |

| | wt | pai/wt | pai/pai | wt | pai/wt | pai/pai |
|---|---|---|---|---|---|---|
| 20:5n-3 | 0.01 ± 0.00 | 0.01 ± 0.00 | 0.01 ± 0.00 | 0.03 ± 0.01 | 0.03 ± 0.01 | 0.02 ± 0.01 |
| 24:1 | 0.05 ± 0.01 | 0.06 ± 0.01 | 0.06 ± 0.01 | 0.05 ± 0.01 | 0.05 ± 0.01 | 0.05 ± 0.01 |
| 22:5n-3 | 0.17 ± 0.04 | 0.16 ± 0.03 | 0.17 ± 0.03 | 0.18 ± 0.33 | 0.10 ± 0.04 | 0.08 ± 0.04 |
| 22:6n-3 | 2.46 ± 0.40 | 2.37 ± 0.40 | 2.74 ± 0.48 | 1.24 ± 0.49 | 1.67 ± 0.68 | 1.60 ± 0.29 |
| Total FAs | 10.59 ± 0.88 | 10.83 ± 0.90 | 11.54 ± 1.11 | 9.59 ± 0.99[a] | 12.97 ± 2.21[b] | 10.66 ± 1.08[c] |
| SFAs | 3.29 ± 0.30[a] | 3.29 ± 0.38[a] | 3.69 ± 0.33[b] | 3.59 ± 0.40[a] | 4.57 ± 0.91[b] | 4.01 ± 0.39[b] |
| Unsaturated FAs | 7.30 ± 0.59 | 7.20 ± 0.91 | 7.85 ± 0.79 | 6.00 ± 0.63[a] | 7.74 ± 1.68[b] | 6.65 ± 0.69[ab] |
| MuFAs | 1.19 ± 0.15[a] | 1.27 ± 0.17[ab] | 1.44 ± 0.15[b] | 1.20 ± 0.19[a] | 1.83 ± 0.39[b] | 1.37 ± 0.15[c] |
| PUFAs | 6.09 ± 0.49 | 5.92 ± 0.75 | 6.40 ± 0.69 | 4.77 ± 0.52[a] | 5.88 ± 1.51[b] | 5.26 ± 0.57[ab] |
| n-6 | 3.45 ± 0.54 | 3.37 ± 0.46 | 3.47 ± 0.30 | 3.32 ± 0.35[a] | 4.07 ± 0.91[b] | 3.55 ± 0.33[ab] |
| n-3 | 2.64 ± 0.39 | 2.55 ± 0.41 | 2.93 ± 0.49 | 1.45 ± 0.31[a] | 1.81 ± 0.73[b] | 1.71 ± 0.33[a] |
| n-6/n-3 | 1.35 ± 0.36 | 1.34 ± 0.20 | 1.21 ± 0.20 | 2.37 ± 0.48 | 2.43 ± 0.64 | 2.13 ± 0.37 |
| n7/(n7+16:0) | 0.20 ± 0.02 | 0.21 ± 0.01 | 0.20 ± 0.01 | 0.16 ± 0.01[a] | 0.19 ± 0.02[b] | 0.16 ± 0.01[a] |
| n7/(n7+18:0) | 0.29 ± 0.02 | 0.30 ± 0.02 | 0.31 ± 0.03 | 0.32 ± 0.03[a] | 0.37 ± 0.07[b] | 0.33 ± 0.01[a] |
| (n-6-LA)/n-6 | 0.40 ± 0.03 | 0.40 ± 0.02 | 0.39 ± 0.02 | 0.66 ± 0.05 | 0.66 ± 0.03 | 0.66 ± 0.04 |
| ***Livers*** | | | | ***Skeletal muscle*** | | |
| No. of samples | 12 | 11 | 10 | 9 | 8 | 8 |
| 8:0 | 0.01 ± 0.00 | 0.01 ± 0.00 | 0.01 ± 0.01 | 0.03 ± 0.01 | 0.04 ± 0.02 | 0.03 ± 0.01 |
| 10:0 | ND | ND | ND | ND | ND | ND |
| 12:0 | ND | ND | ND | 0.02 ± 0.01 | 0.01 ± 0.01 | 0.01 ± 0.00 |
| 14:0 | 0.02 ± 0.00 | 0.02 ± 0.01 | 0.02 ± 0.01 | 0.03 ± 0.01 | 0.04 ± 0.02 | 0.10 ± 0.20 |
| 14:1 | 0.01 ± 0.00 | 0.01 ± 0.00 | 0.01 ± 0.00 | ND | ND | ND |

| | wt | pai/wt | pai/pai | wt | pai/wt | pai/pai |
|---|---|---|---|---|---|---|
| 16:0 | 1.95 ± 0.34 | 1.89 ± 0.34 | 1.77 ± 0.40 | 1.04 ± 0.26 | 1.03 ± 0.36 | 0.97 ± 0.35 |
| 16:1n-7 | 0.14 ± 0.08 | 0.14 ± 0.05 | 0.12 ± 0.06 | 0.12 ± 0.03 | 0.15 ± 0.07 | 0.10 ± 0.03 |
| 18:0 | 1.52 ± 0.24[a] | 1.51 ± 0.39[ab] | 1.35 ± 0.41[b] | 0.61 ± 0.16 | 0.58 ± 0.22 | 0.60 ± 0.25 |
| trans-18:1 | 0.02 ± 0.01 | 0.02 ± 0.02 | 0.02 ± 0.01 | ND | ND | ND |
| 18:1n-9 | 1.29 ± 0.36 | 1.27 ± 0.26 | 1.21 ± 0.27 | 0.35 ± 0.10 | 0.43 ± 0.21 | 0.34 ± 0.15 |
| 18:1n-7 | 0.28 ± 0.13[a] | 0.31 ± 0.06[a] | 0.22 ± 0.09[b] | 0.14 ± 0.04 | 0.14 ± 0.05 | 0.15 ± 0.07 |
| 18:2n-6 | 1.29 ± 0.20[a] | 1.09 ± 0.25[b] | 1.07 ± 0.23[b] | 0.87 ± 0.41 | 0.84 ± 0.31 | 0.61 ± 0.17 |
| 20:0 | 0.01 ± 0.01 | 0.02 ± 0.01 | 0.01 ± 0.01 | ND | ND | ND |
| 18:3n-6 | 0.02 ± 0.00[a] | 0.02 ± 0.00[ab] | 0.02 ± 0.00[b] | ND | ND | ND |
| 18:3n-3 | 0.01 ± 0.00 | 0.01 ± 0.00 | 0.01 ± 0.00 | ND | ND | ND |
| t10c12-CLA | 0.012 ± 0.01 | 0.017 ± 0.01 | 0.010 ± 0.00 | ND | ND | ND |
| t10c12-CLA (%) | 0.13 ± 0.09 | 0.19 ± 0.12 | 0.14 ± 0.05 | ND | ND | ND |
| 20:3n-6 | 0.20 ± 0.06[a] | 0.18 ± 0.05[ab] | 0.15 ± 0.04[b] | ND | ND | ND |
| 20:4n-6 | 1.46 ± 0.32 | 1.50 ± 0.41 | 1.41 ± 0.52 | 0.57 ± 0.16 | 0.52 ± 0.19 | 0.54 ± 0.32 |
| 20:5n-3 | 0.03 ± 0.01 | 0.02 ± 0.01 | 0.02 ± 0.01 | ND | ND | ND |
| 24:1 | 0.02 ± 0.01 | 0.03 ± 0.01 | 0.02 ± 0.00 | 0.06 ± 0.01 | 0.24 ± 0.13 | 0.27 ± 0.19 |
| 22:5n-3 | 0.04 ± 0.01 | 0.03 ± 0.01 | 0.03 ± 0.01 | 0.08 ± 0.02 | 0.08 ± 0.05 | 0.08 ± 0.03 |
| 22:6n-3 | 0.85 ± 0.18 | 0.86 ± 0.17 | 0.73 ± 0.28 | 0.76 ± 0.29 | 0.71 ± 0.21 | 0.75 ± 0.19 |
| Total FAs | 9.19 ± 1.55[a] | 8.94 ± 1.59[ab] | 8.19 ± 1.78[b] | 4.66 ± 1.34 | 4.69 ± 1.45 | 4.42 ± 1.41 |
| SFAs | 3.51 ± 0.47 | 3.45 ± 0.61 | 3.16 ± 0.71 | 1.74 ± 0.44 | 1.69 ± 0.60 | 1.70 ± 0.56 |
| Unsaturated FAs | 5.67 ± 1.12[a] | 5.49 ± 0.99[ab] | 5.03 ± 1.10[b] | 2.92 ± 0.92 | 3.00 ± 0.87 | 2.72 ± 0.87 |
| MuFAs | 1.76 ± 0.53 | 1.78 ± 0.33 | 1.59 ± 0.35 | 0.64 ± 0.14 | 0.84 ± 0.39 | 0.73 ± 0.34 |

| | wt | pai/wt | pai/pai | wt | pai/wt | pai/pai |
|---|---|---|---|---|---|---|
| PUFAs | 3.90 ± 0.67[a] | 3.70 ± 0.75[ab] | 3.44 ± 1.01[b] | 2.28 ± 0.82 | 2.16 ± 0.54 | 1.99 ± 0.61 |
| n-6 | 2.97 ± 0.54[a] | 2.78 ± 0.61[ab] | 2.65 ± 0.74[b] | 1.43 ± 0.54 | 1.36 ± 0.42 | 1.16 ± 0.47 |
| n-3 | 0.93 ± 0.19 | 0.92 ± 0.17 | 0.79 ± 0.29 | 0.85 ± 0.31 | 0.79 ± 0.24 | 0.83 ± 0.21 |
| n-6/n-3 | 3.28 ± 0.62 | 3.02 ± 0.36 | 3.59 ± 0.80 | 1.69 ± 0.39 | 1.87 ± 0.75 | 1.40 ± 0.44 |
| n7/(n7+16:0) | 0.17 ± 0.05 | 0.19 ± 0.03 | 0.16 ± 0.05 | 0.20 ± 0.02 | 0.22 ± 0.01 | 0.20 ± 0.03 |
| n7/(n7+18:0) | 0.45 ± 0.07 | 0.46 ± 0.04 | 0.48 ± 0.11 | 0.36 ± 0.04 | 0.42 ± 0.07 | 0.36 ± 0.03 |
| (n-6-LA)/n-6 | 0.56 ± 0.04 | 0.60 ± 0.05 | 0.58 ± 0.08 | 0.41 ± 0.08 | 0.39 ± 0.10 | 0.45 ± 0.07 |
| ***BAT*** | | | | ***WAT*** | | |
| No. of samples | 8 | 8 | 7 | 5 | 3 | 3 |
| 8:0 | 0.01 ± 0.00 | 0.01 ± 0.00 | 0.01 ± 0.00 | ND | ND | ND |
| 10:0 | 0.01 ± 0.00 | 0.01 ± 0.00 | 0.01 ± 0.00 | ND | ND | ND |
| 12:0 | 0.03 ± 0.05 | 0.01 ± 0.00 | 0.01 ± 0.00 | 0.007 ± 0.00 | 0.005 ± 0.00 | 0.008 ± 0.00 |
| 14:0 | 0.21 ± 0.07 | 0.20 ± 0.04 | 0.20 ± 0.05 | 0.040 ± 0.02[ab] | 0.031 ± 0.01[a] | 0.051 ± 0.00[b] |
| 14:1 | 0.01 ± 0.00 | 0.01 ± 0.00 | 0.01 ± 0.00 | 0.006 ± 0.00 | 0.005 ± 0.00 | 0.006 ± 0.00 |
| 16:0 | 2.85 ± 0.57 | 2.78 ± 0.44 | 2.95 ± 1.10 | 0.592 ± 0.27[ab] | 0.529 ± 0.03[a] | 0.684 ± 0.03[b] |
| 16:1n-7 | 0.67 ± 0.31 | 0.55 ± 0.17 | 0.65 ± 0.16 | 0.186 ± 0.07 | 0.170 ± 0.01 | 0.189 ± 0.01 |
| 18:0 | 1.30 ± 0.21 | 1.29 ± 0.31 | 1.28 ± 0.36 | 0.155 ± 0.07[ab] | 0.132 ± 0.02[a] | 0.165 ± 0.01[b] |
| trans-18:1 | 0.10 ± 0.04 | 0.08 ± 0.05 | 0.09 ± 0.05 | 0.013 ± 0.00 | 0.017 ± 0.01 | 0.022 ± 0.01 |
| 18:1n-9 | 3.47 ± 0.78 | 3.08 ± 0.84 | 3.25 ± 0.98 | 0.843 ± 0.44[ab] | 0.807 ± 0.03[a] | 0.936 ± 0.01[b] |
| 18:1n-7 | 0.80 ± 0.27 | 0.68 ± 0.16 | 0.75 ± 0.22 | 0.108 ± 0.03[a] | 0.118 ± 0.00[a] | 0.162 ± 0.02[b] |
| 18:2n-6 | 1.88 ± 0.31 | 1.70 ± 0.42 | 1.69 ± 0.31 | 0.504 ± 0.34[ab] | 0.333 ± 0.01[a] | 0.438 ± 0.01[b] |
| 20:0 | 0.03 ± 0.00 | 0.03 ± 0.01 | 0.03 ± 0.01 | ND | ND | ND |

| | wt | pai/wt | pai/pai | wt | pai/wt | pai/pai |
|---|---|---|---|---|---|---|
| 18:3n-6 | 0.03 ± 0.01 | 0.02 ± 0.00 | 0.02 ± 0.01 | 0.007 ± 0.00 | 0.006 ± 0.00 | 0.004 ± 0.00 |
| 18:3n-3 | 0.04 ± 0.01 | 0.03 ± 0.01 | 0.04 ± 0.01 | 0.022 ± 0.01[ab] | 0.014 ± 0.00[a] | 0.019 ± 0.00[b] |
| t10c12-CLA | 0.026 ± 0.02 | 0.026 ± 0.01 | 0.029 ± 0.01 | ND | 0.004 ± 0.00 | 0.005 ± 0.00 |
| t10c12-CLA (%) | 0.20 ± 0.13 | 0.22 ± 0.08 | 0.23 ± 0.07 | ND | 0.130 | 0.128 |
| 20:3n-6 | 0.07 ± 0.01 | 0.07 ± 0.01 | 0.10 ± 0.09 | 0.010 ± 0.00 | 0.008 ± 0.00 | 0.011 ± 0.00 |
| 20:4n-6 | 1.11 ± 0.24 | 1.10 ± 0.27 | 1.01 ± 0.19 | 0.081 ± 0.02 | 0.081 ± 0.02 | 0.087 ± 0.00 |
| 20:5n-3 | 0.02 ± 0.01 | 0.02 ± 0.01 | 0.02 ± 0.00 | ND | ND | ND |
| 24:1 | 0.02 ± 0.01 | 0.02 ± 0.01 | 0.03 ± 0.01 | ND | ND | ND |
| 22:5n-3 | 0.02 ± 0.00 | 0.03 ± 0.01 | 0.02 ± 0.00 | 0.006 ± 0.00 | 0.005 ± 0.00 | 0.006 ± 0.00 |
| 22:6n-3 | 0.29 ± 0.07 | 0.32 ± 0.07 | 0.30 ± 0.07 | 0.035 ± 0.02 | 0.022 ± 0.01 | 0.027 ± 0.00 |
| Total FAs | 12.99 ± 2.07 | 12.02 ± 2.41 | 12.48 ± 3.30 | 3.373 ± 1.52 | 3.087 ± 0.25 | 3.893 ± 0.49 |
| SFAs | 4.43 ± 0.64 | 4.31 ± 0.74 | 4.48 ± 1.49 | 0.796 ± 0.36[ab] | 0.698 ± 0.04[a] | 0.910 ± 0.04[b] |
| Unsaturated FAs | 8.55 ± 1.52 | 7.71 ± 1.75 | 8.00 ± 1.86 | 1.895 ± 0.87[ab] | 1.665 ± 0.02[a] | 2.003 ± 0.05[b] |
| MuFAs | 5.07 ± 1.27 | 4.41 ± 1.16 | 4.78 ± 1.35 | 1.185 ± 0.54[ab] | 1.157 ± 0.03[a] | 1.362 ± 0.04[b] |
| PUFAs | 3.45 ± 0.53 | 3.28 ± 0.74 | 3.19 ± 0.54 | 0.680 ± 0.38[ab] | 0.487 ± 0.03[a] | 0.615 ± 0.02[b] |
| n-6 | 3.07 ± 0.52 | 2.88 ± 0.68 | 2.82 ± 0.48 | 0.608 ± 0.35[ab] | 0.439 ± 0.02[a] | 0.556 ± 0.02[b] |
| n-3 | 0.38 ± 0.06 | 0.40 ± 0.07 | 0.38 ± 0.08 | 0.066 ± 0.03 | 0.042 ± 0.01 | 0.052 ± 0.00 |
| n-6/n-3 | 8.22 ± 1.74 | 7.21 ± 1.09 | 7.62 ± 1.07 | 8.948 ± 1.23[a] | 10.556 ± 1.59[ab] | 10.707 ± 0.71[b] |
| n7/(n7+16:0) | 0.33 ± 0.06 | 0.30 ± 0.05 | 0.33 ± 0.05 | 0.342 ± 0.06[ab] | 0.353 ± 0.00[a] | 0.339 ± 0.01[b] |
| n7/(n7+18:0) | 0.72 ± 0.05 | 0.70 ± 0.08 | 0.72 ± 0.03 | 0.838 ± 0.02 | 0.859 ± 0.02 | 0.850 ± 0.01 |

| | wt | pai/wt | pai/pai | wt | pai/wt | pai/pai |
|---|---|---|---|---|---|---|
| (n-6-LA)/n-6 | 0.39 ± 0.03 | 0.41 ± 0.03 | 0.40 ± 0.02 | 0.213 ± 0.10 | 0.240 ± 0.03 | 0.212 ± 0.03 |

Note: Tissues of wt, pai/wt, and pai/pai mice at age of 11 weeks are for analysis, and 19:0 is an internal standard. Each value represents the mean ± SD. ND, Not detected; LA, linoleic acid; c9t11, cis 9, trans 11-CLA; FA, fatty acids. Total FAs are total amounts from 8:0 to 22:6n-3 compositions. SFAs (saturated) and MUFAs (monounsaturated) are calculated as 8:0 + 10:0 + 12:0 + 14:0 + 16:0 + 18:0 + 20:0 + 22:0 + 24:0, and 14:1 + 16:1n-7 + trans-18:1 + 18:1n-9 + 18:1n-7 + 20:1 + 22:1 + 24:1, respectively. PUFAs is calculated as n-6 + n-3 as well as n-6 and n-3 are calculated as 18:2n-6 + 18:3n-6 + 20:3n-6 + 20:4n-6, and 18:3n-3 + 20:5n-3 + 22:5n-3 + 22:6n-3, respectively. n-7/(n-7 + 16:0) and n-7/(n-7 + 18:0) are calculated as (16:1n-7 + 18:1n-7)/(16:1n-7 + 18:1n-7 + 16:0), and 18:1n-9/(18:1n-9 + 18:0), respectively. Different letters in superscript indicate a level of statistical significance of $p < 0.05$ within the same gender.

**Supplementary Table 5. Dissection parameters in wt and pai mice**

| | wt | pai/wt | pai/pai |
|---|---|---|---|
| Number of mice | 21 | 16 | 14 |
| Bodyweight (g) | 26.8 ± 1.7 | 27.9 ± 1.6 | 26.8 ± 1.3 |
| Heart weight (%) | 0.47 ± 0.03 | 0.48 ± 0.12 | 0.57 ± 0.18 |
| Liver weight (%) | 4.60 ± 0.57[a] | 4.88 ± 0.83[a] | 5.91 ± 0.96[b] |
| Spleen weight (%) | 0.34 ± 0.07[a] | 0.38 ± 0.17[ab] | 0.48 ± 0.20[b] |
| Kidney weight (%) | 1.24 ± 0.08[a] | 1.31 ± 0.18[ab] | 1.44 ± 0.23[b] |
| Lung weight (%) | 0.60 ± 0.18 | 0.59 ± 0.18 | 0.66 ± 0.23 |
| Pancreas weight (%) | 0.51 ± 0.07 | 0.52 ± 0.16 | 0.54 ± 0.18 |
| Brain weight (%) | 1.18 ± 0.10 | 1.20 ± 0.22 | 1.32 ± 0.34 |
| Hypothalamus weight (%) | 0.03 ± 0.01 | 0.03 ± 0.01 | 0.03 ± 0.01 |
| Testis weight (%) | 0.59 ± 0.13 | 0.58 ± 0.20 | 0.59 ± 0.24 |
| WAT weight (%) | 1.69 ± 0.47[a] | 1.88 ± 0.64[a] | 1.35 ± 0.46[b] |
| BAT weight (%) | 0.39 ± 0.07[a] | 0.50 ± 0.18[ab] | 0.54 ± 0.18[b] |
| Small intestine length (cm) | 36.5 ± 2.6 | 35.6 ± 2.2 | 36.5 ± 3.0 |

Note: Mice at 11 weeks of age were killed in the non-fasted state. Tissue/organ weight as a percentage of slaughter bodyweight was calculated. BAT, interscapular brown adipose tissue; WAT, epididymal white adipose tissue. Each value represents the mean ± SD. Different letters in superscript indicate $p < 0.05$ within the same gender.
